# Nonspecific Membrane Interactions Can Modulate BK Channel Activation

**DOI:** 10.1101/2020.02.05.936161

**Authors:** Mahdieh Yazdani, Guohui Zhang, Zhiguang Jia, Jingyi Shi, Jianmin Cui, Jianhan Chen

## Abstract

Large-conductance potassium (BK) channels are transmembrane (TM) proteins that can be synergistically and independently activated by membrane voltage and intracellular Ca^2+^. The only covalent connection between the cytosolic Ca^2+^ sensing domain and the TM pore and voltage sensing domains is a 15-residue “C-linker”. To determine the linker’s role in BK activation, we designed a series of linker sequence scrambling mutants to suppress potential complex interplay of specific interactions with the rest of the protein. The results revealed a surprising sensitivity of BK activation to the linker sequence. Combing atomistic simulations and further mutagenesis experiments, we demonstrated that nonspecific interactions of the linker with membrane alone could directly modulate BK activation. The C-linker thus plays more direct roles in mediating allosteric coupling between BK domains than previously assumed. Our results also suggest that covalent linkers could directly modulate TM protein function and should be considered an integral component of the sensing apparatus.

**One-sentence summary:** The covalent linker between BK intracellular and transmembrane domains is likely a part of the sensing apparatus that regulates the channel activation.

## Introduction

Widely distributed in nerve and muscle cells, large-conductance potassium (BK) channels are characterized by a large single-channel conductance (∼ 100 - 300 pS) (1-5) and dual activation by both intracellular Ca^2+^ and membrane voltage (6-8), thus an ideal model system for understanding the gating and sensor-pore coupling in ion channels. BK channels are involved in numerous vital physiological processes including intracellular ion homeostasis and membrane excitation, and are associated with pathogenesis of many diseases such as epilepsy, stroke, autism and hypertension (9). Functional BK channels are homo-tetramers, each containing three distinct domains (Figure 1a). The voltage sensor domain (VSD) detects membrane potential, the pore-gate domain (PGD) controls the K^+^ selectivity and permeation, and the cytosolic tail domain (CTD) senses various intracellular ligands including Ca^2+^. The VSD and the CTD also form a Mg^2+^ binding site for Mg^2+^ dependent activation (Yang 2008 NSMB). The tetrameric assembly of CTD domains is also referred to as the “gating ring”. VSD and PGD together form the trans-membrane domain (TMD) of BK channels. Previous studies on mouse BK channels (10) and recent atomistic structures of full-length *Aplysia californica* BK channel (aSlo1) (11, 12) reveal that CTD of each subunit reside beneath the TMD of the neighboring subunit in a surprising domain-swapped arrangement (Figure S1a).

**Figure 1.**
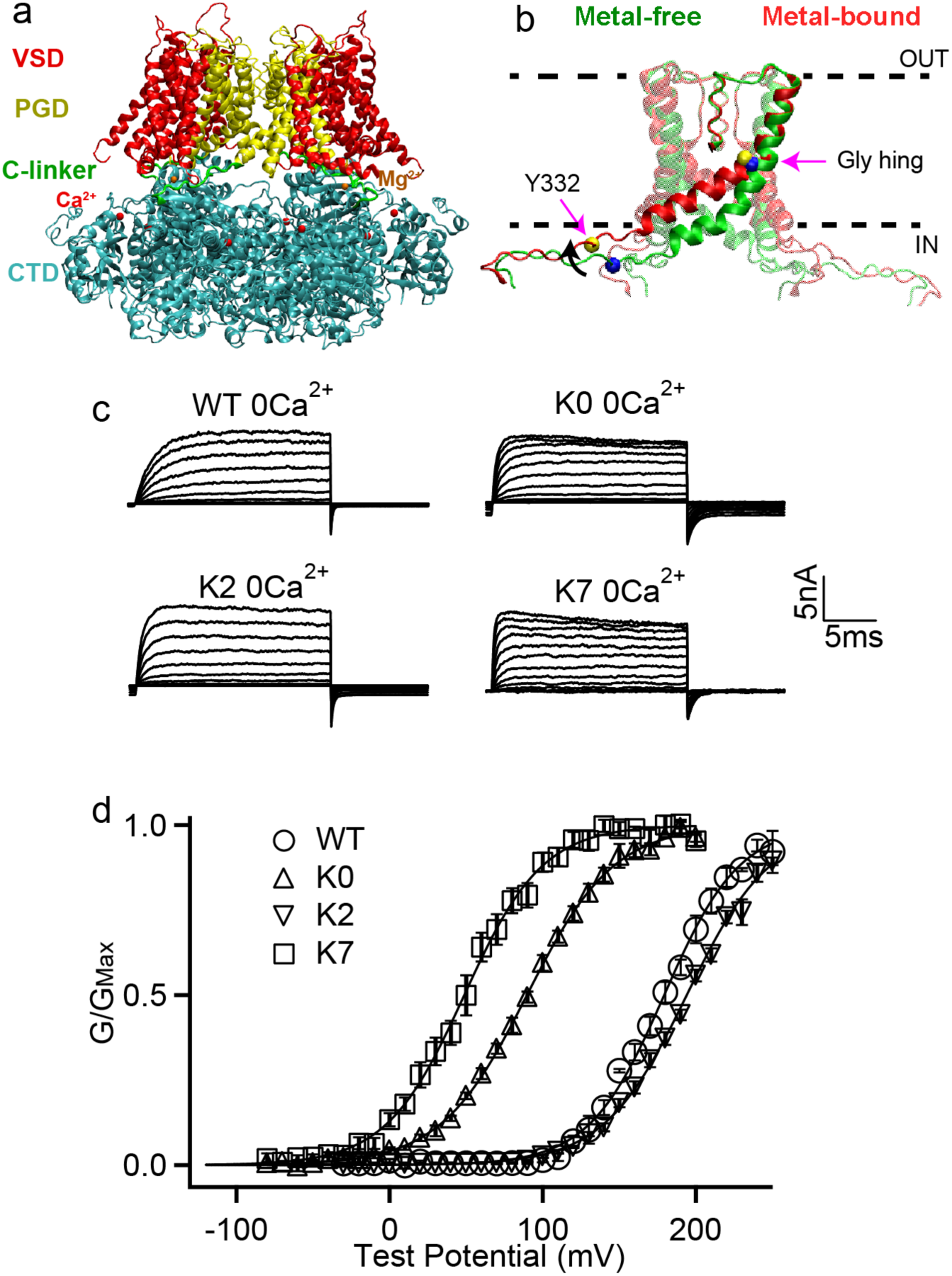
Structure of BK channels and effects of C-linker sequence scrambling on its voltage activation **a.** Key functional domains and structural organization of the human BK channel (hSol1). See main text for definition of domains; **b**. Orientation of the pore lining S6 helices and C-linkers in the metal-free (green) and metal-bound (red) states; The dash lines indicate the approximate positions of membrane interfaces. The location of Tyr 332 and Gly 310 are indicated by yellow and blue spheres in the metal-bound and metal-free states, respectively. The black arrow shows S6 movement upon metal binding. **c.** Macroscopic currents of WT, K0, K2 and K7 hSlo1 channels. The currents were elicited in 0 [Ca^2+^]_i_ by voltage pulses from −30 to 250 mV with 20 mV increments for WT and K2 and voltage pulses from −80 to 200 mV with 20 mV increments for K0 and K7. The voltages before and after the pulses were −50 and −80 mV, respectively. **d.** Conductance-voltage (G-V) curves for WT, K0, K2 and K7 hSlo1 channels in 0 [Ca^2+^]_i_ showing significant shifts in the activation voltage (V_0.5_); All solid lines were fit to the Boltzmann relation (see Methods), with V_0.5_ of 183.4 ± 3.2 mV for WT; 89.6 ± 3.5 mV for K0; 195.5 ± 3.5 mV for K2; and 48.7 ± 4.7 mV for K7.

The only covalent connection between CTD and TMD of BK channels is a 15-residue peptide referred to as the “C-linker” (R329 to K343 in the human BK channel, hSlo1) (green in Figure 1a). This linker directly connects the pore lining S6 helices in the PGD (yellow in Figure S1b) to the N-terminus of CTD (known as RCK1 N-lobe; blue in Figure S1b), and is believed to play an important role in mediating the gating ring-pore coupling (1, 8, 9). For example, *Niu* et al. (13) observed that lengthening the C-linker through inserting poly-AAG (Table S1) was accompanied with right shifted G-V curves, while shortening the C-linker led to left-shifted G-V curves, clearly demonstrating the importance of C-linker in BK gating. Intriguingly, the voltage required for half activation, V_0.5_, displayed a highly linear relationship with the number of residues inserted or deleted in the absence of Ca^2+^. This led to the proposal that the linker-gating ring behaves as a “passive spring” in activation of BK channels (13), suggesting that the C-linker largely provides an inert linkage.

Evidence has also accumulated to suggest that the C-linker may play more direct roles in mediating allosteric coupling of BK channels. Both Ca^2+^-free and bound Cryo-EM structures of full-length aSlo1 (11, 12) were found to contain wide-open pores and thus did not provide clue to how the channel might be gated. Atomistic simulations later revealed that, due to the movement of pore-lining S6 helices (Figure 1b), the BK pore cavity become narrower, more elongated, and crucially, more hydrophobic in the metal-free state (i.e., without bound Ca^2+^ and Mg^2+^; presumably the closed state) (14). As such, the pore can readily undergo hydrophobic dewetting transition to give rise to a vapor barrier that prevents ion permeation (14). Recognition of hydrophobic gating in BK channels provides a mechanistic basis for further understanding how the C-linker may mediate the sensor-pore coupling. Specifically, key movements of S6 helices involve bending at the glycine hinge (G310, G311) toward the membrane upon Ca^2+^ binding (Figure 1b), leading to ∼6 Å expansion of the pore entrance near I323. Considering that C-linkers are directly connected to S6 helices (Figure S1b), their interactions with the rest of the channel as well as membrane and water will likely have an effect on the S6 orientation and consequently BK channel activation. It has also been proposed that BK channels may be gated at the selectivity filter (15-17), the conformation of which could be modulated by the S6 helix orientation and thus C-linker interactions. The notion that interactions of C-linkers can modulate BK activation has actually been demonstrated in a recent study, where the R_329_KK_331_ residues in the C-linker was proposed to form alternating interactions with E321 and E224 from the neighboring chains and membrane lipids during each gating cycle (18).

A key challenge of using mutagenesis to delineate the roles of a specific residue or interaction in protein function is that multiple competing effects may be perturbed simultaneously. The C-linker, in particular, is involved in extensive interactions with the gating ring and VSD (Figure S2), making it very difficult to derive unambiguous interpretation of its role in BK activation (18). In this work, we examine a set of BK channels with scrambled C-linker sequences to determine if the C-linker is largely inert in mediating the sensor-pore coupling of BK channels. Combining atomistic simulation, mutagenesis and electrophysiology, we discover that the C-linker is a major pathway of gating ring-pore communication and can play a much more direct and specific role in mediating BK activation than previously thought. In particular, we show that nonspecific interactions of the C-linker with the membrane/solvent environment, namely, membrane anchoring effects, can directly modulate voltage gating of BK channels. This observation has been verified by additional experiments performed on BK constructs either lacking CTD or with the membrane anchoring residue mutated.

## Results and Discussions

### Sequence scrambling of the C-linker dramatically modulates BK voltage activation

To further resolve whether the C-linker is largely an inert linkage (as part of the linker-gating ring “passive spring”) or it has more direct roles (mediated through various specific and nonspecific interactions of C-linker residues), we investigated a series of BK mutants where the C-linker sequence is scrambled (Table 1). If the linker-gating ring largely acts like a passive spring with an inert C-linker, the expectation is that these scrambling mutant BK channels would have similar gating properties. Table 1 shows experimental results on the scramble mutant channels. Among these mutants K3, K5 and K6 did not show functional expression, while other mutant channels showed robust currents (Figure 1c). We measured voltage dependent activation of these channels and the conductance-voltage (G-V) relationships were fitted using the Boltzmann function to derive *V*_0.5_, the voltage where *G*/*G*_Max_ reaches 0.5 (Figure 1d). Left shift of G-V (*V*_0.5_ decreases) indicates that the channel requires less voltage to activate, while right shift of G-V (*V*_0.5_ increases) indicates that the channel is opened by higher voltage (See Methods for details). Importantly, as shown in Figure 1d and Table 1, V_0.5_ depends very sensitively on the C-linker sequence. All five mutants that lead to functional channels, K0, K1, K2, K4 and K7, have significantly altered activation voltage, with V_0.5_ changes as large as ∼135 mV. This is in contrast to the prediction of a largely inert C-linker.

**Table 1:** C-linker scrambling mutations and measured V_0.5_ in the full-length and Core-MT BK channels at [Ca^2+^] = 0 and [Mg^2+^] = 0. The Core-MT constructs are based on the TMD, C-linker of mSlo1, and an 11-residue tail from K_V_ 1.4 of the mouse Shaker family (19, 20). The location of the nearest Tyr to the S6 C-terminal is highlighted in green. K0 (Y330G) was designed to remove the Tyr sidechain in the K0 background.

| Mutation | Sequence | | | | | | $V_{0.5}$ (mV) | |
| --- | --- | --- | --- | --- | --- | --- | --- | --- |
|  |  |  |  |  |  |  | Full-Length | Core-MT |
| WT | EIIEL | IGNRK | K<u>Y</u>GGS | YSAVS | GRK |  | 183.4 | 235.0 |
| K0 | EIIEL | IGNR<u>Y</u> | GKGSK | YSRAV | SKG |  | 89.6 | 192.6 |
| K0(Y330G) | EIIEL | IGNR<u>G</u> | GKGSK | YSRAV | SKG |  | 169.8 | NM |
| K1 | EIIEL | RIGNK | <u>Y</u>GGSY | KSAVR | KSG |  | 136.2 | NM |
| K2 | EIIEL | IGRKN | <u>Y</u>KGGS | YSARV | SGK |  | 195.5 | 263.5 |
| K3 | EIIEL | IGN<u>Y</u>G | GRSYS | KAKVS | RKG |  | NC | NM |
| K4 | EIIEL | IGRN<u>Y</u> | GGSYS | AKKVR | SKG |  | 94.6 | NM |
| K5 | EIIER | LIGKK | RN<u>Y</u>KG | GSYSA | VSG |  | NC | NM |
| K6 | EIIEL | RKKIR | KGN<u>Y</u>G | GSYSA | VSG |  | NC | NM |
| K7 | EIIEL | IGN<u>Y</u>G | GSYSA | VRKSK | GRK |  | 48.7 | 167.5 |

NC: no current, channel could not be expressed; NM: Mutation Not Made

### C-linker is a structured loop with limited flexibility and a key dynamic pathway of BK gating ring-pore coupling

Atomistic modeling and simulation were performed to investigate possible mechanisms of unexpected sensitivity of BK gating on the C-linker sequence. For this, we first derived atomistic structures of the wild-type (WT) hSlo1 based on the aSlo1 Cryo-EM structures (11, 12) (see Methods). The homology models have been cross validated using Cryo-EM structures of hSlo1 that became available after the completion of this work (21). The results show that aS1o1-derived models are essentially identical to the actual hSlo1 structures, with the central PGD structures differ by only 0.87 Å (Figure S2). Atomistic simulations also suggest that the homology models and Cryo-EM structures lead to similar structural and dynamical properties (e.g., see Figure S3). These structures reveal that the C-linker is a structured loop with essentially identical conformations in both metal-free (closed) and metal-bound (open) states (Figure 1b). The C_α_ root-mean-square deviation (RMSD) of C-linker conformations between the two states is only ∼0.8 Å. The linker forms extensive contacts with the RCK1 N-lobe (H344 to N427) of the gating ring, mainly mediated by the Y_332_GGSYSA_338_ segment in the C-linker, and S0’ of VSD (Figure S3). In particular, the two conserved tyrosine residues (Y332 and Y336) are fully embedded in a hydrophobic RCK1 N-lobe pocket, apparently maintaining a tight packing between the C-linker and RCK1 N-lobe (Figure S3). Several positively charged residues (R329, K330, K331, R342 and K343) flank the above segment and are exposed to solvent, likely mediating the solubility of the C-linker. Importantly, these two short tracks of charged residues appear to be tightly anchored by the C-linker-RCK1 contacts.

The stability of the C-linker conformation as a structured loop is further confirmed by atomistic molecular dynamic (MD) simulations, which showed that the C-linker maintained stable conformations and contacts with VSD and RCK1 N-lobe throughout the 800 ns simulation time (Figures 2a-b and S5-8, WT). The root-mean-square fluctuation (RMSF) of the C-linker region is ∼1-2 Å, which is only slightly elevated compared to regions with helical or sheet secondary structures (Figure 2b). The end-to-end distance of the C-linker fluctuated stably ∼30 Å in both metal-bound and free state simulations (Figure 2a), even as the pore underwent dewetting transitions in the metal-free state (Figure S8c).

**Figure 2.**
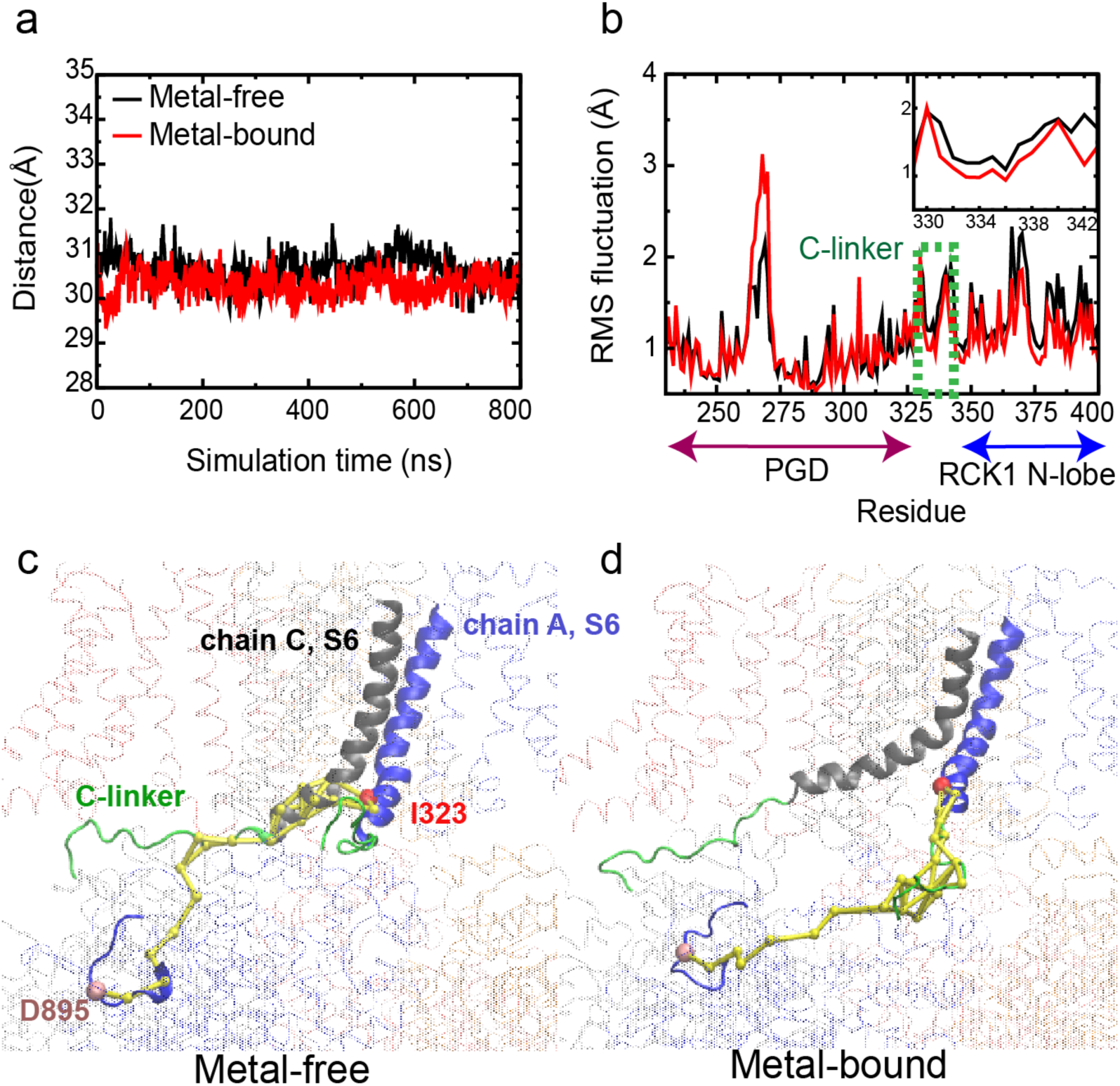
Dynamic properties of BK channels. **a.** The C-linker N-C Cα distance during a representative 800-ns MD simulations of the WT hSlo1. **b.** Residue Root-Mean-Square Fluctuations (RMSF) profiles of PGD, C-linker and RCK1 N-lobe derived from the same trajectory, showing limited flexibility of the C-linker (green dashed box and shown in the insert). **c-d.** Optimal and top 10 suboptimal pathways of dynamic coupling (yellow traces) between D895 in the RCK2 Ca^2+^ binding site of chain A (C_α_ colored as pink) and I323 in PGD of chain A (C_α_ colored as red) in the metal-free (e) and bound (f) states. The C-linker is colored green, S6 of chain A and chain C in blue and grey, respectively. The rest of the channel is shown as transparent ribbons. Note that the pathways in the metal-free state go through the neighboring chain (chain C).

Dynamic network analysis further reveals that the C-linker is a key pathway of dynamic coupling between the gating ring and PGD. Such analysis utilizes correlation of residue motions during MD simulations to uncover probable pathways of allosteric coupling in biomolecules (22-24). The optimal and suboptimal paths of coupling are then identified as the shortest paths with the highest pairwise correlations, which should possess the highest probabilities of information transfer (22). We analyzed the optimal and suboptimal pathways of coupling between I323 in the S6 helices, where substantial conformational change occurs during the gating event (14), and critical metal binding residues, including D895 and R514 in the RCK2 and RCK1 Ca^2+^ binding site, respectively, and E374 in the Mg^2+^ binding site residing in the CTD. The results reveal that the main pathways of communication from Ca^2+^ and Mg^2+^ binding sites to the PGD, go through the C-linker for all four chains (Figure 2c-d and S9). This is not necessarily surprising since C-linker is the only covalent connection between the domains. Interestingly though, in the metal-free state the main path from the RCK2 Ca^2+^ binding site to PGD goes from the neighboring monomer in every other chain (Figure 2c-d). This can be attributed to much tighter S6 helix packing in the metal-free state (14), and, combined with the domain swapped arrangement of BK tetramers, may help enforce cooperative gating response upon metal binding.

### Scrambling mutations minimally perturb C-linker structure and dynamics

Atomistic simulations suggest that the changes in voltage dependent gating are unlikely to derive from a change of the structural features or functional roles of the permutated linkers. The structure of the channel appears minimally perturbed by the mutated linkers, with the overall RMSD below ∼5 Å and the TM domain (core) RMSD around 2-3 Å from the initial cryo-EM-derived structures for both WT and mutant channels (Figure S8a-b). All mutant channels can readily undergo hydrophobic dewetting transitions as observed for the WT channel (Figure S8c). The linker region also maintains similar backbone conformations in WT and all mutants (Figure S7), even though it becomes slightly more dynamic in the K7 mutant as reflected in the RMSF profile (Figure S7b). Furthermore, scrambling mutations do not appear to perturb long-range coupling properties either; the C-linker remains to provide the key pathway of dynamic coupling between CTD and PGD.

Sequence properties, particularly distributions of charged residues, can modulate intrinsic conformational properties of disordered peptides including chain extension (25), which could explain the changes in channel activation in terms of the linker-gating ring passive spring model. To investigate this hypothesis, we performed atomistic simulations of the isolated C-linker segments in the ABSINTH implicit solvent (26), which has been shown to reliably predict the inherent conformational extension of disordered peptides (27). The results suggest that V_0.5_ of linker scrambling mutants is not correlated with the inherent linker extension (Figure S10) and is inconsistent with the prediction based on the previous linker-gating ring passive spring model (13) (dashed line in Figure S10). This observation further supports the notion that the C-linker is unlikely an inert component of the linker-gating ring passive spring.

Finally, since the C-linker is a structured loop with extensive interactions with the CTD and VSD (Figure S2, S5-6), we considered whether some specific subsets of mutations could perturb the coupling among VSD, PGD and CTD to modulate voltage dependent activation (V_0.5_), even though sequence scrambling is designed to suppress such effects. To test this, we analyzed C-linker residue contact probability maps from MD trajectories to identify potential group of contacts that could be correlated to V_0.5_ (Figure S5-6). However, none of the observed changes in specific protein-protein contacts could explain the measured V_0.5_ variations.

### Tyr membrane anchoring effects can modulate S6 orientation to affect BK activation

As discussed above, key conformational changes involved in activation of BK channels include the re-orientation of the pore-lining S6 helices, which bends outwardly and toward the membrane at the glycine hinge (G310 and G311) (Figure 1b). Directly connected to the S6 helix, C-linker residues would be moved closer to the membrane interface during activation (e.g., comparing red vs green cartoons in Figure 1b). As such, it can be anticipated that nonspecific interactions of the C-linker with the membrane interface can affect channel activation by stabilizing the bended conformation of S6 helices, thus modulating the activation voltage. Among various amino acids, aromatic ones such as Tyr and Trp are known to be “membrane anchoring”, with a strong preference towards localizing at the membrane-water interface (28, 29). While the sequence scrambling was designed to suppress the potential effects of specific interactions of the C-linker, the two Tyr residues (Y332 and Y336 in WT) were placed at different separations from the membrane interface (Table 1, highlighted in green fonts). Indeed, the measured V_0.5_ shows a strong correlation with the sequence separation between E324 (the approximate location of membrane interface) and the nearest C-linker Tyr residue (Figure 3). Positioning this Tyr residue closer to the membrane interface could allow stronger membrane anchoring, preferentially stabilizing the bended conformation of S6 in the open state and shifting the equilibrium towards the active state of the channel as observed.

**Figure 3.**
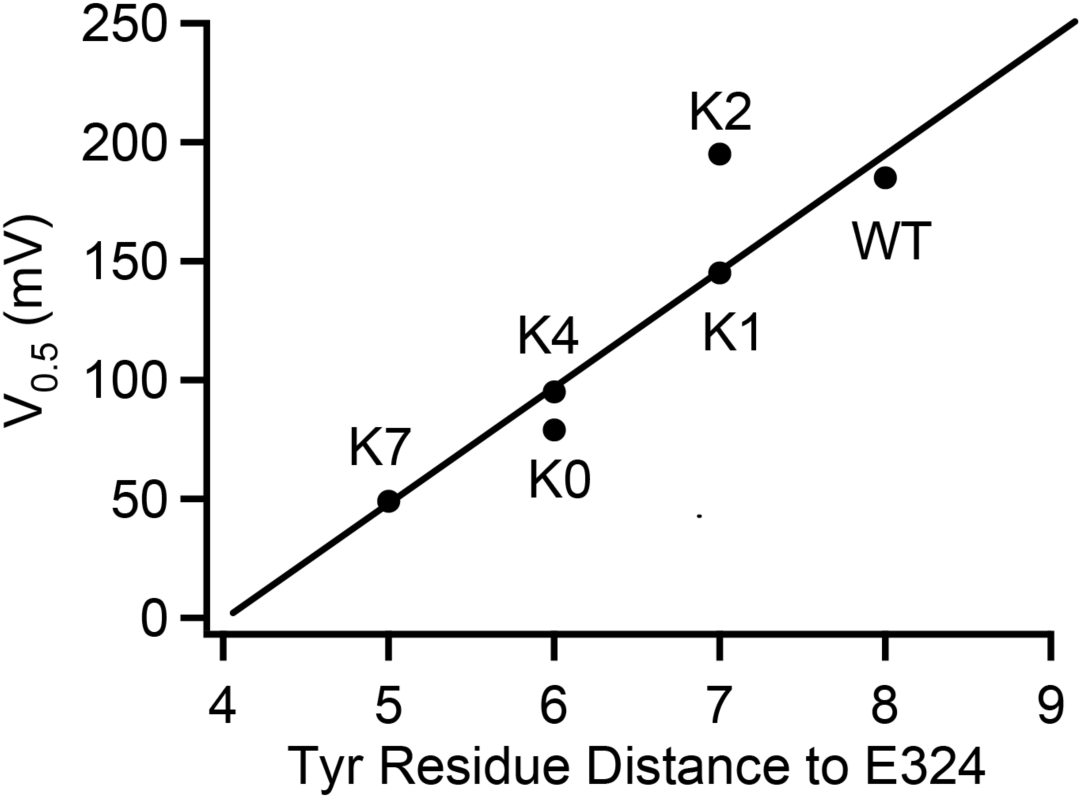
Correlation between V_0.5_ and the sequential distance (residue counts) of the nearest C-linker Tyr (Y332 in WT; Y330 in K0; Y331 in K1; Y331 in K2; Y330 in K4; Y329 in K7) to the S6 C-terminal (E324) in the C-linker scrambling mutants.

To more directly examine if membrane anchoring plays a role in modulating BK activation, we further analyzed the atomistic trajectories to understand the details of Tyr interaction with the membrane interface. As illustrated in Figure 4, Tyr sidechains contain both aromatic rings and polar groups that allow them to embed their aromatic rings in the lipid tail region and at the same time direct the polar -OH groups toward the lipid headgroup region to form hydrogen bonding interactions with water and lipid headgroups (29). In addition, the aromatic ring could also form π-cation interactions with positively charged cholines in lipid headgroups (30). Importantly, results from simulation analysis confirm that positioning of the C-linker in the metal-bound state allows more extensive interactions between Try sidechains and the membrane interface (e.g., compare Figure 4 a vs b and c vs d), which would stabilize the open state and thus lower the voltage required for activation. This is more clearly shown in the average hydrophobic, hydrogen-bonding and π-cation interactions formed by the nearest Tyr sidechains (Y332 in WT, Y330 in K0, Y331 in K2, and Y329 in K7; see Table 1), summarized in Figures 5 and S11. This is consistent with the observation that Tyr membrane anchoring lowers V_0.5_ significantly for all mutants except K2. Note that Y329 of K7 appears to be dominated by extensive π-cation interactions in the metal-bound state (Figure 5d) and as a consequence forms slightly fewer hydrogen bonding interactions on average (Figure 5b). Consistent with our experimental data (Figure 1 c-d), placing Tyr closer to the end of the S6 helix indeed allows more extensive Tyr/membrane interface interactions, forming larger numbers of polar and nonpolar interactions on average (Figure 5). However, we note that directly quantifying the free energy contribution of Tyr membrane anchoring effects is technically difficult. It requires rigorous calculation of the free energy of the open/close transition of the whole channel, which is not yet feasible given the current computational capability. In particular, there is substantial heterogeneity in the lipid distribution near the channel, especially around the C-linker region (e.g., see Figures 4 and S12). Achieving convergence on the free energy of activation transition would require sufficient sampling of these lipid configurations, which has been shown to be extremely slow at the multi-μs level or longer (31).

**Figure 4.**
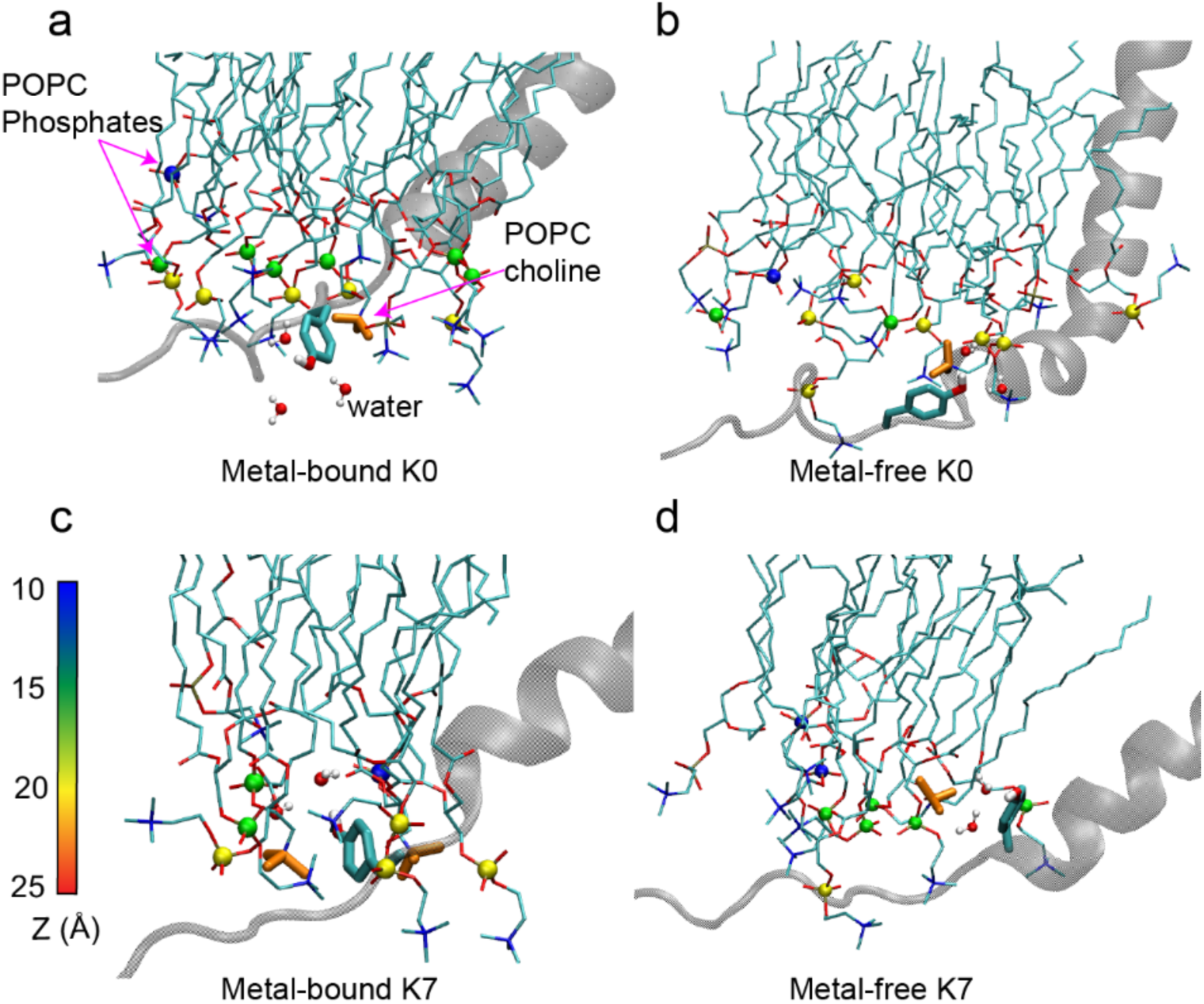
Interactions of the C-linker Tyr residue nearest to the S6 helix with the membrane interface in hSlo1K0 (Y330) and K7 (Y329) mutants. Only the S6 helix and C-linker from one subunit are shown for clarity. POPC molecules near the Tyr residue are shown in sticks, with the phosphorous atoms shown in spheres and colored according to their distance to the membrane center (Z). POPC choline groups (orange Licorice) and water molecule near the Tyr sidechain are also shown to illustrate the π-cation and hydrogen bonding interactions. Note that the C-linker is positioned closer to the interface and forms more extensive interactions in the metal-bound (activated) state.

**Figure 5.**
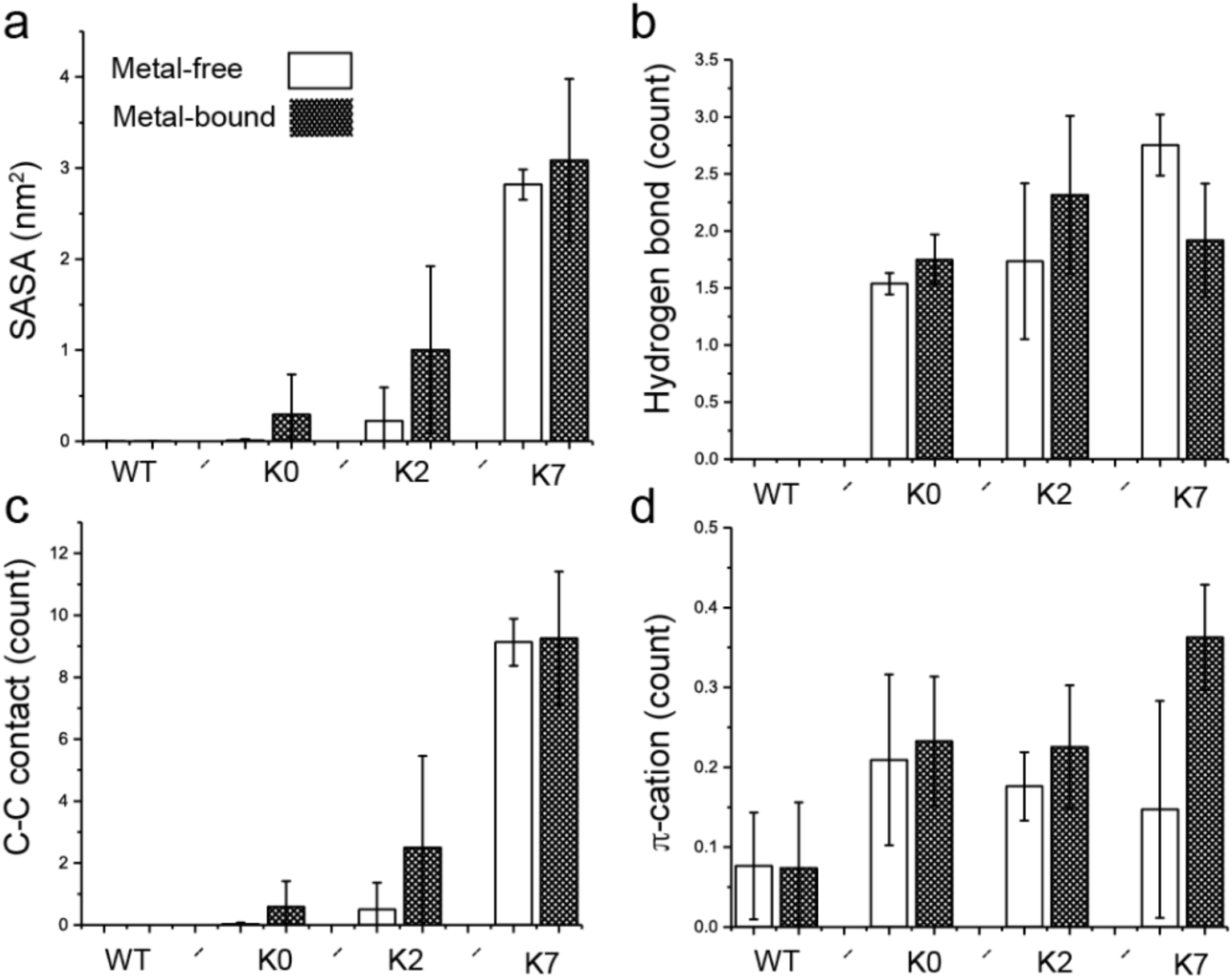
Interactions of the C-linker Tyr residue nearest to the S6 helix with the membrane interface. **a.** Average Tyr sidechain solvent accessible surface area (SASA) of burial by lipid tails, representing the level of hydrophobic contacts between the Tyr sidechain and aliphatic lipid tails. **b.** Average number of hydrogen bonds between the Tyr OH group and the POPC hydrophilic head groups. **c.** Average number of carbon-carbon (C-C) contacts between the Tyr aromatic ring and POPC hydrocarbon tails. **d.** Average number of π-cation interactions between the Tyr aromatic ring and POPC choline group. All results are the average of three independent simulations, with the standard error shown as the error bar. No hydrophobic, hydrogen bonding, or C-C contacts as defined above were observed in the WT channel.

### Tyr membrane anchoring affects BK channel activation similarly without the gating ring

If the observed effects on the voltage gating of full-length BK channels are indeed mainly due to the nonspecific interactions between the C-linker and membrane interface, they should persist in the Core-MT truncated channel (19, 20), in which the whole gating-ring is removed and there is no CTD coupling with either VSD or PGD via the C-linker (32) (Fig 5a). To test this prediction, Core-MT channels with three C-linker scrambling mutations, K0, K2 and K7, were expressed (Fig 5b) and their voltage dependent activation measured. Mutations of the C-linker shifted the G-V relation of the Core-MT constructs in the same directions as observed for the full-length channels (Table 1), with K0 and K7 stabilizing channel activation (Fig 5c, e) while K2 making activation at higher voltages (Fig 5d). However, the reduction in the activation voltage by K0 mutation is only ∼42 mV in Core-MT, compared to ∼94 mV in the full-length channel (Figure 6c), and by K7 mutation is ∼67 mV in Core-MT, compared to ∼135 mV in the full-length channel (Figure 6e). This is not considered surprising as the C-linker is more flexible in the absence of the gating ring, thus weakening the effects of membrane anchoring. The observation that linker sequence scrambling mutations can modulate BK activation even in the Core-MT background is remarkable, providing a direct evidence that the C-linker is more than an inert, passive covalent linker for coupling the gating ring and PGD. Instead, the linker plays a more direct and more specific role in modulating the opening of BK pore, such as through its interactions with the membrane environment. As such, the linker could be considered an integral component of the sensing apparatus.

**Figure 6.**
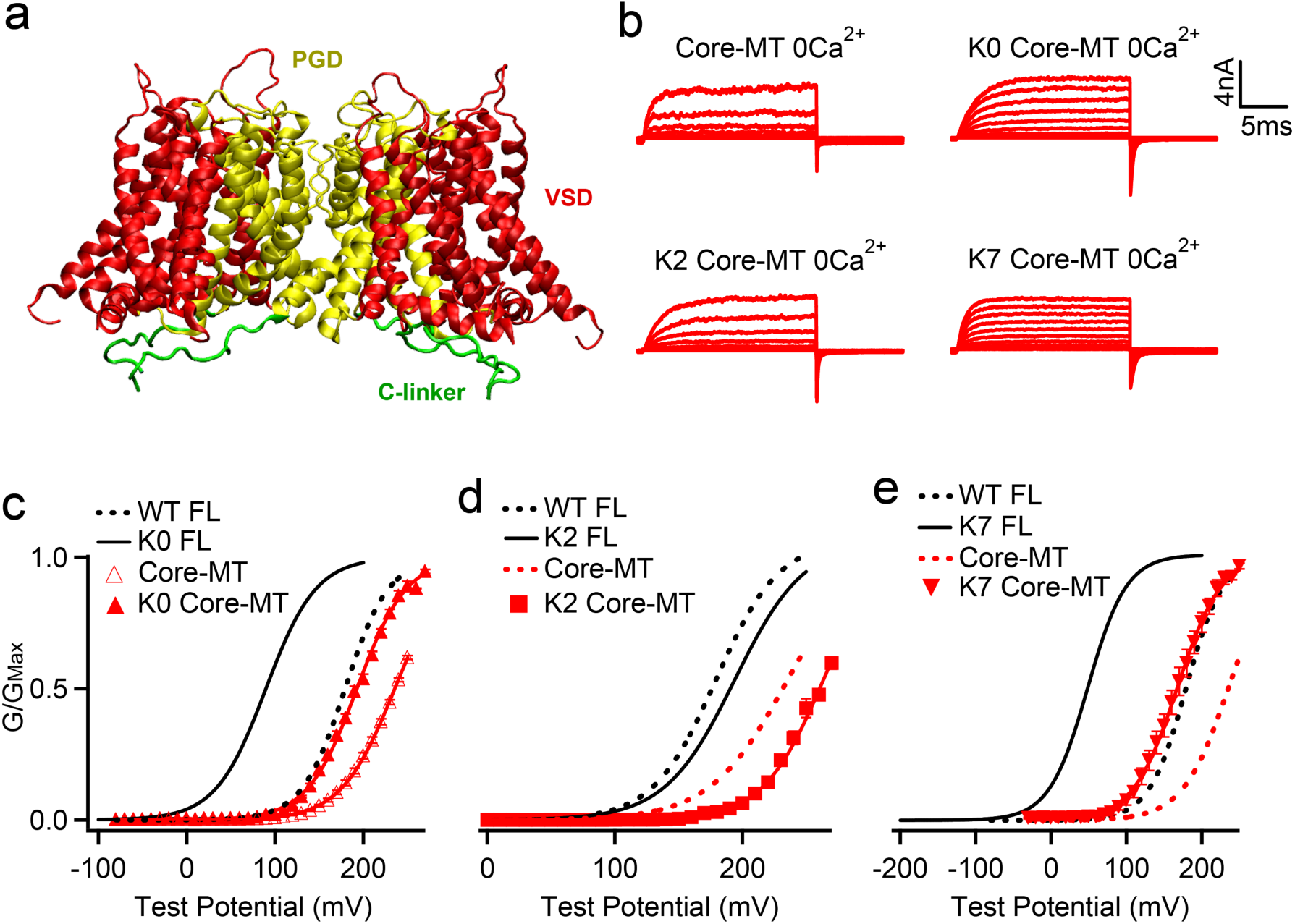
Effects of C-linker scrambling mutations of voltage activation of Core-MT BK channels. **a.** Illustration of the structure of the Core-MT BK channel, where the whole CTD is absent. The mini-tail is omitted in the illustration for clarity. **b.** Macroscopic currents of the WT Core-MT construct as well as the K0, K2 and K7 C-linker scrambling mutants. The currents were elicited in 0 [Ca^2+^]_i_ by voltage pulses from - 30 to 250 mV with 20 mV increments. The voltages before and after the pulses were −50 and −80 mV, respectively. **c-e.** G-V curves for WT Core-MT and K0, K2 and K7 mutants in 0 [Ca^2+^]_i_ (red traces) The G-V curves of the Full-Length (FL) channels are also shown for reference (black traces). All lines were fit to the Boltzmann relation (see Methods) with V_0.5_ of 235.0 ± 3.1 mV for WT Core-MT; 192.6 ± 3.8 mV for K0 Core-MT; 263.5 ± 4.0 mV for K2 Core-MT; and 167.5 ± 4.2 mV for K7 Core-MT.

### Removal of membrane anchoring Tyr in a C-linker BK mutant recovers WT-like gating

We note that Tyr is not the only type of residues being shuffled in the sequence scrambling (Table 1) and that interactions of other residues, particularly charged ones (18), with membrane and water could also affect the open/close equilibrium of the channel. This may explain why K1 and K2 mutant channels have different V_0.5_, even though the nearest Tyr is at position 331 in both mutants (Table 1). To further examine if Tyr residues indeed provide the dominant contributions, we replaced Y330 with Gly in the K0 mutant to completely remove the aromatic side chain. The mutant expressed robust currents (Figure 7a). Strikingly, K0 Y330G mutation abolished the effects of K0 on G-V relation and largely shifted the G-V back to that of the WT, with a V_0.5_ of 169.8 ± 5.0 mV as compared to 183.4 ± 3.2 mV for the WT (Figure 7b). This, together with the correlation shown in Figure 3, provides a direct support that membrane anchoring effects of Tyr are mainly responsible for modulating the activation voltage in the C-linker sequence scrambling mutants.

**Figure 7.**
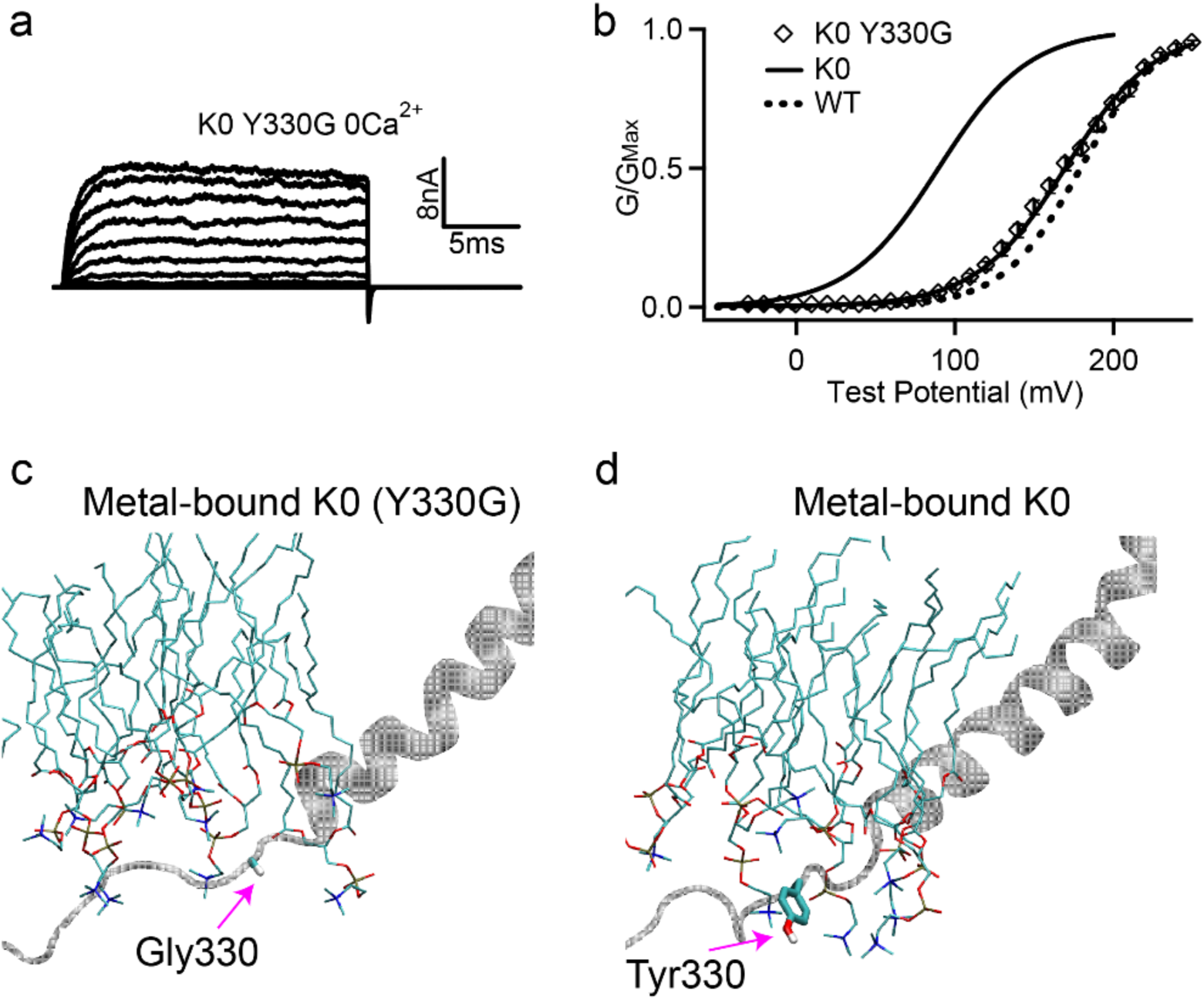
Removing Tyr sidechain in the K0 mutant full-length hSlo1 channel recovers WT-like voltage activation. **a.** Macroscopic currents of K0 (Y330G) mutant channel. The currents were elicited in 0 [Ca^2+^]_i_ by voltage pulses from −30 to 250 mV with 20 mV increments. The voltages before and after the pulses were −50 and −80 mV, respectively. **b.** The G-V curve in 0 [Ca^2+^]_i_, showing the recovery of activation voltage (V_0.5_) of the K0 (Y330G) to the WT level. The solid line was fit to the Boltzmann relation (see Methods) with 169.8 ± 5.0 mV for the K0 (Y330G). The G-V curves of the full-length WT and K0 mutant channels are also shown for reference (solid and dash lines). **c-d**: Representative molecular snapshots showing the effects of Y3330G mutation in K0 channels. See Figure 4 caption for the details of molecular representations.

## Conclusion

We have combined atomistic simulations and experiment to determine the role of the 15-residue C-linker in the sensor-pore coupling of BK channels. As the only covalent connection between PGD and CTD of BK channels, the C-linker has been widely assumed to play an important role in mediating the information transfer and domain coupling. Our analysis show that C-linker is a structured loop with extensive contacts with CTD and VSD, and remain highly stable in both metal-bound (activated) and metal-free (deactivated) states. Dynamic coupling analysis confirms that C-linker is the key pathway of the senor-pore coupling. However, the linker is not just an inert pathway. Instead, sequence scrambling of the C-linker can greatly affect the activation voltage of hSlo1 channels. Atomistic simulations revealed that the effects on the activation voltage could be mainly attributed to the nonspecific interactions between the C-linker and membrane interface, particularly membrane anchoring effects of Tyr residues. These conclusions were supported by additional experiments showing that the effects of C-linker sequence scrambling persist in the Core-MT constructs, where the gating ring is completely removed, and that replacing the key Tyr residue with Gly largely abolishes the shift in the G-V curve in a selected C-linker mutant. To the best of our knowledge, this is one of the first direct demonstrations of how nonspecific membrane interactions can modulate TM protein function.

Our conclusion that the C-linker is unlikely an inert component of the linker-gating ring “passive spring” is also consistent with other structural and functional studies of BK channels. For example, the Cryo-EM structures of full-length BK channels (11, 12) reveal that the previously published poly-AAG insertion site (13), right after residue S337 (Table S1), actually locates in a short loop following the C-linker segment -Y_332_GGSY_336_-that forms stable contacts with RCK1 N-lobe (Figure S2). The inserted residues would project away from the channel (Figure S13), and are very unlikely to affect the effective C-linker length (or the gating ring-pore distance) as originally designed. Instead, the observed effects in BK gating V_0.5_ upon insertion/deletion of C-linker residues could likely be attributed to certain nontrivial structural and/or dynamical impacts, such as weakening of the VSD/CTD interactions. Another important evidence that is inconsistent with the passive spring model comes from the study of Core-MT BK channels. Since the whole gating ring is removed, the Core-MT construct should correspond to a state where the passive spring is fully relaxed and thus V_0.5_ maximizes. Yet, the V_0.5_ of WT Core-MT is only ∼52 mV larger than the full-length BK channel (Table 1). This is far below what may be expected based on poly-AAG insertion mutants, which can increase V_0.5_ by ∼142 mV with (AAG)_3_ inserted (13) (also see Table S1). Interestingly, the recently published Cryo-EM structures confirms that β subunits make extensive contacts with the C-linker for influencing the channel gating and manipulating the channel function (21).

TM ion channels and receptors frequently contain separate TM domains, which directly mediate function such as ion permeation, and intracellular and extracellular sensing domains, which often control the function of TM domains in response to various chemical signals (33). The general role of the covalent linkers connecting the sensing and TM domains is of great general interest (34, 35). A central question is whether the linker mainly provides an inert and passive connection between sensing and TM domains or it should be considered an element of signal sensing itself. Dissecting the potential roles of covalent linkers is challenging because multiple sources of interactions and conformational transitions could impose competing strains on the linker to modulate functional regulation. It is difficult to design experiments that can unambiguously test and validate whether a particular type of strain imposed on the covalent linker could lead to predictable functional outputs. Linker sequence permutation could provide an effective strategy to suppress the potential consequence of specific (but unknown) linker interactions, allowing one to test the functional role of a single type of strain imposed on the linker (such as membrane anchoring). Our findings show that non-specific interactions of the C-linker can regulate BK voltage gating. Therefore, covalent linkers of membrane proteins could serve as sensors of signals that perturb their interaction with the environment, which in term can modulate the functional center in the TM domain, may it be the gate of ion channels or intramolecular signaling pathway in receptors.

## Acknowledgements

We thank Rohit Pappu for the original design of C-linker mutants. This work was supported by National Institutes of Health Grants R01 HL70393 and GM114300 (to Chen). We also would like to thank Roderick MacKinnon and Xiao Tao for sharing the hSlo1 Cryo-EM structures prior to their release in PDB. Computing for this project was performed on the Pikes cluster housed in the Massachusetts Green High-Performance Computing Center (MGHPCC).

## Materials and Methods

### Homology modeling and atomistic simulations

As described previously (14), homology models of the WT metal-bound and metal-free hSol1 channels were built using Modeller v9.14 (36) based on sequence alignment of Tao et al (11). Sequence alignment shows 55.95% identity for the full-length channels. The sequence identities in PGD is higher at 61.96%. The high level of sequence identity suggests that the homology models are likely reliable. This has been confirmed by direct comparison with the recently published Cryo-EM structures of hSlo1 (21) (Figures S2). The backbone RMSD between the model and new structure is only 2.18 Å at the whole channel level and as low as 0.87 Å in the PGD. Structures for C-linker scrambling mutants were build based on the WT hSlo1 models using CHARMM (37).

Using CHARMM-GUI server (38), the hSlo1 structures were inserted in POPC lipid bilayers followed by solvation using the TIP3P water model (39). 450 mM KCl was then added, same as used in Cryo-EM structure determination (11, 12). Each system was first energy minimized, followed by multiple cycles of equilibration dynamics with gradually decreasing harmonic restrains on positions of selected protein/lipid heavy atoms. To ensure that the size of the simulation box become stable, in the last equilibration step, only protein heavy atoms were harmonically restrained while the system equilibrated under NPT (constant particle number, pressure and temperature) condition. The final simulation box size is ∼18 x 18 x 15.4 nm^3^ with ∼476,000 atoms, containing ∼97,000 water molecules and ∼800 lipid molecules. The Charmm36m all-atom force field (40) was used for all systems. The production simulations were performed using CUDA-enabled Amber14 (41). The MD time step was set at 2 fs. Particle Mesh Ewald (PME) algorithm (42) with a cut-off at 12 Å was used to describe the electrostatic interactions. Van der Waals interactions were cutoff at 12 Å with a smooth switching function starting from 10 Å. Lengths of all covalent bonds involving hydrogen atoms were fixed using SHAKE algorithm (43). The system temperature was maintained at 298 K by Langevin dynamics with a friction coefficient of 1 ps^-1^, and the pressure was maintained semi-isotopically at 1 bar at both x and y (membrane lateral) directions using the Monte-Carlo barostat method (44, 45). Three independent 800-ns NPT production simulations were performed for each construct (WT, K0, K2 and K7) in both metal-free and bound states, with an aggregated simulation time of 19.2 μs. Snapshots were saved every 50 ps for post analysis.

### Structural and dynamic analysis

All analysis were performed using a combination of in-house scripts, MDAnalysis (46) and Gromacs2016 (47, 48) software. Only snapshots from the last 150 ns of all production MD trajectories were used for the calculation of Tyr-membrane interactions (SASA of burial, hydrogen bonding, π-cation and carbon-carbon contacts) as well as RMSF and the C-linker contact map (SI). A (hydrophobic) carbon-carbon contact was considered formed if the distance is no greater than 4.5 Å. The π-cation interaction was identified when the distance between the center of mass of the Tyr aromatic ring and Nitrogen atom of POPC choline group is no greater than 5.0 Å. Similarly, the cutoff was set at 5.0 Å for calculation of the C-linker contact map (SI) while only the heavy atoms of each residue were considered. Hydrogen bonds were analyzed using the MDAnalysis “HyrogenBondAnalysis” class with default criteria. The number of pore water molecules was calculated using the same criteria as described previously (14). Dynamic network analysis was performed using the *Networkview* (22) plugin of VMD (49). For this, snapshots were extract every 1 ns from the 800 ns molecular dynamic trajectories. To build the network, each amino acid was represented as a single node at their Cα position and a contact (edge) was defined between two nodes if the minimal heavy-atom distance between them was within a cutoff distance (4.5 Å) during at least 75% of the trajectory. The resulting contact matrix were then weighted based on the correlation coefficients of dynamic fluctuation (*<u>C</u>*_ij_), calculated using the Carma software (50), as w_ij_ = −log (|C_ij_|), where C_ij_ = <Δr_i_(t).Δr_j_(t)> / (<Δr_i_(t)^2^> <Δr_j_(t)^2^>)^1/2^ and Δr_i_(t) = r_i_(t) - <r_i_(t)>, r_i_(t) is the position of the atom corresponding of the i^th^ node and <> denotes ensemble average (over the MD trajectory). The path length between the desired nodes were then calculated as the sum of the edge weights. The shortest (optimal) path, calculated using Floyd-Warshall algorithm (51), is believed to represent the dominant mode of communication. Slightly longer (suboptimal) paths were also calculated. VMD was used for preparing all molecular illustrations.

### Mutations and Expression

Mutations in all experiments were made by using overlap-extension PCR (polymerase chain reaction) with Pfu polymerase (Stratagene) from the mbr5 splice variant of *mslo1* (Uniprot ID: Q08460) (52). And then all PCR-amplified regions were verified by sequencing (53). RNA was then transcribed in vitro from linearized DNA with T3 polymerase (Ambion, Austin, TX) and an amount of 0.05–50 ng/oocyte RNA was injected into oocytes (stage IV-V) from female *Xenopus laevis*. After injection, these oocytes were incubated at 18°C for 2-7 days.

### Electrophysiology

We used inside-out patches to record Ionic currents with an Axopatch 200-B patch-clamp amplifier (Molecular Devices, Sunnyvale, CA) and ITC-18 interface and Pulse acquisition software (HEKA Elektronik GmbH, Holliston, MA). Borosilicate pipettes those were used for inside-out patches had 0.5–1.5 MΩ resistance and then form patches from oocyte membrane. The current signals were record at 50 KHz sample rate (20-μs intervals) with low-pass-filtered at 10 KHz. In order to remove capacitive transients and leak currents, we applied a P/4 protocol with a holding potential of –120 mV. Our solutions used in recording ionic currents were listed below. **1)** Pipette solution (in mM): 140 potassium methanesulphonic acid, 20 HEPES, 2 KCl, 2 MgCl_2_, pH 7.2. **2)** The nominal 0 µM [Ca^2+^]_i_ solution, which contained about 0.5 nM free [Ca^2+^] _i_ (in mM): 140 potassium methanesulphonic acid, 20 HEPES, 2 KCl, 5 EGTA, and 22 mg/L (+)-18-crown-6-tetracarboxylic acid (18C6TA), pH 7.2. **3)** Basal bath (intracellular) solution (in mM): 140 potassium methanesulphonic acid, 20 HEPES, 2 KCl, 1 EGTA, and 22 mg/L 18C6TA, pH 7.2. Then we added CaCl_2_ into basal solution to obtain the desired free [Ca^2+^]_i_, which was determined by a Ca^2+^-sensitive electrode (Thermo Electron, Beverly, MA). All chemicals were from Sigma-Aldrich unless otherwise noted and all the experiments were done at room temperature (22–24°C).

### Electrophysiology Data analysis

Relative conductance (*G*) was obtained by measuring macroscopic tail current at –80 mV or −120mV. The conductance-voltage (G-V) relationships was plotted to fit with the Boltzmann function:

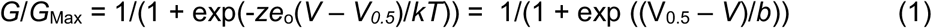

Where *G*/*G*_Max_ means the ratio of conductance to maximal conductance, *z* means the number of equivalent charges, *e*_*o*_ means the elementary charge, *V* means membrane potential, V_0.5_ means the voltage where *G*/*G*_Max_ reaches 0.5, *k* means Boltzmann’s constant, *T* means absolute temperature, and *b* means the slope factor with units of mV. Each G-V relationship was the average of 3–15 patches and error bars in the figures is standard error of means (SEM.).

## Supporting Information

**Table S1.** Effects of C-linker insertion/deletion mutations on the voltage required for half channel activation (V_0.5_) of hSlo1 in absence of Ca^2+^ and Mg^2+^ (data extracted from: Niu et al, 2004). The position of insertion (S337) is colored red.

| Mutation | $V_{0.5}$ (mV) | C-linker Sequence |
| --- | --- | --- |
| -3 | 121 | RKKYGG - - - AVSGRK |
| -1 | 132 | RKKYGGSY- AVSGRK |
| WT | 184 | RKKYGGSY**S**AVSGRK |
| +3 | 210 | RKKYGGSY**S**AAGAVSGRK |
| +6 | 246 | RKKYGGSY**S**AAGAAGAVSGRK |
| +12 | 326 | RKKYGGSY**S**AAGAAGAAGAAGAVSGRK |

**Figure S1.**
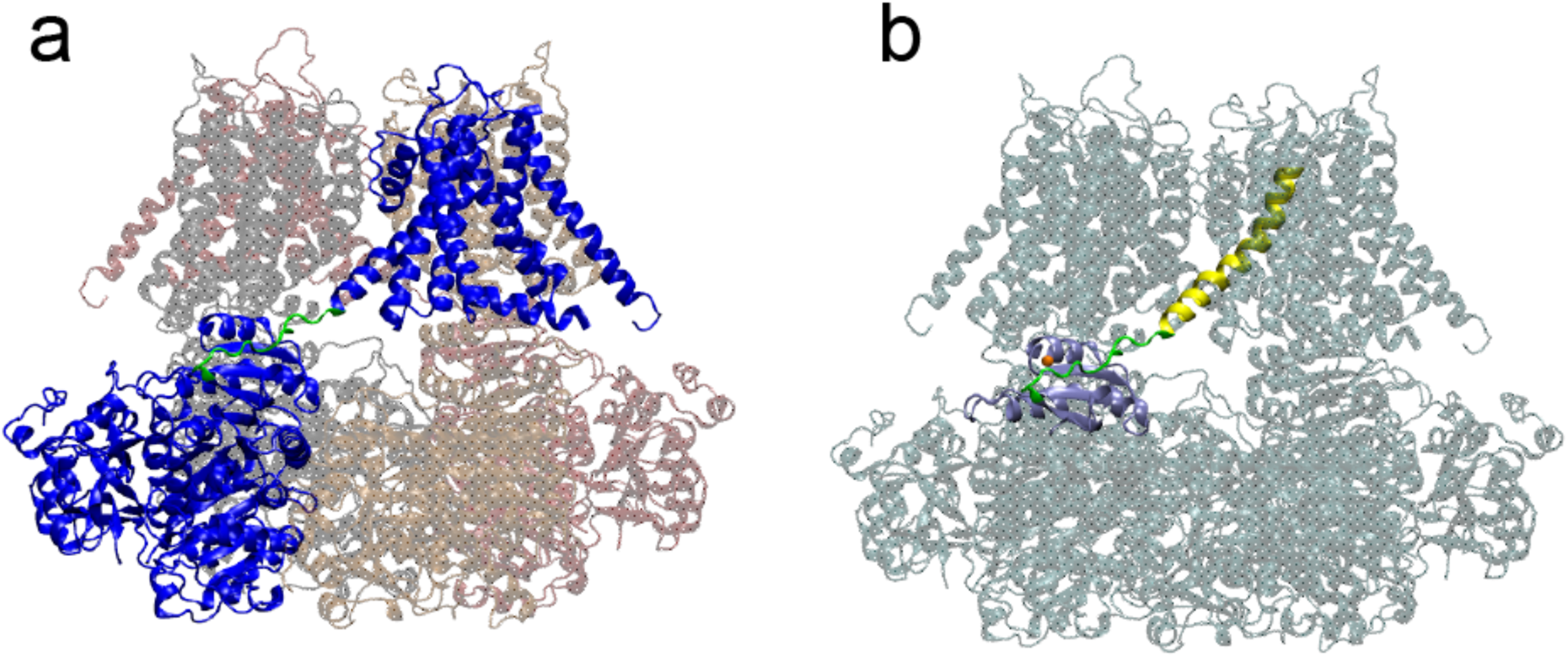
**a.** Domain swapping arrangement of BK channels. Shown is the structure of the human BK channels (hSlo1) with chain A colored blue and C-linker colored green. Other domains are made transparent for clarity. **b.** The C-linker (green) directly connects the pore lining helix (S6, yellow) to the RCK1 N-lobe (blue) of the CTD. The bound Mg^2+^ is shown as an orange sphere.

**Figure S2.**
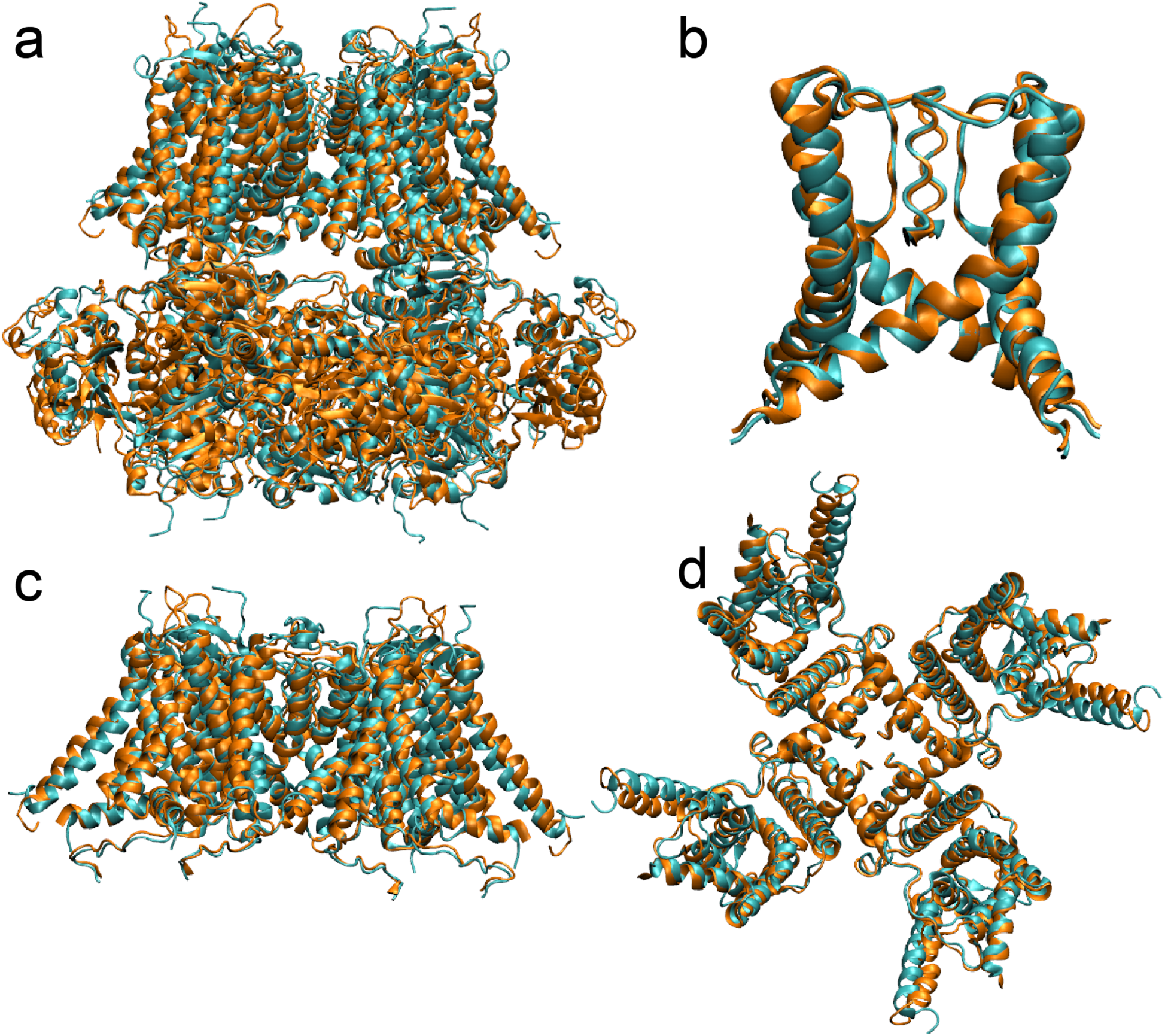
Comparison of the aSlo1-derived homology model of hBK (orange) with the new Cryo-EM structure (PDB 6V38) (cyan) in the metal-bound state. **a.** Overlay of the full-length structures aligned using core residues 100 to 600. The backbone RMSD of two structures 2.18 Å. **b.** Overlay of the central pore region (including S6 helices), aligned based on residues 287 to 330. The backbone RMSD of the central pore is 0.87 Å. **c-d.** Overlay of the whole TMD structures c) side view, d) top view, aligned on residues 100 to 343. The backbone RMSD is 2.53 Å. Note that the main difference is in the relative arrangements of VSD and PGD, which is also observed in multiple Cryo-EM structures of the same aSlo1 or hSlo1 constructs and thus likely reflect inherent flexibility of the VSD/PGD packing.

**Figure S3.**
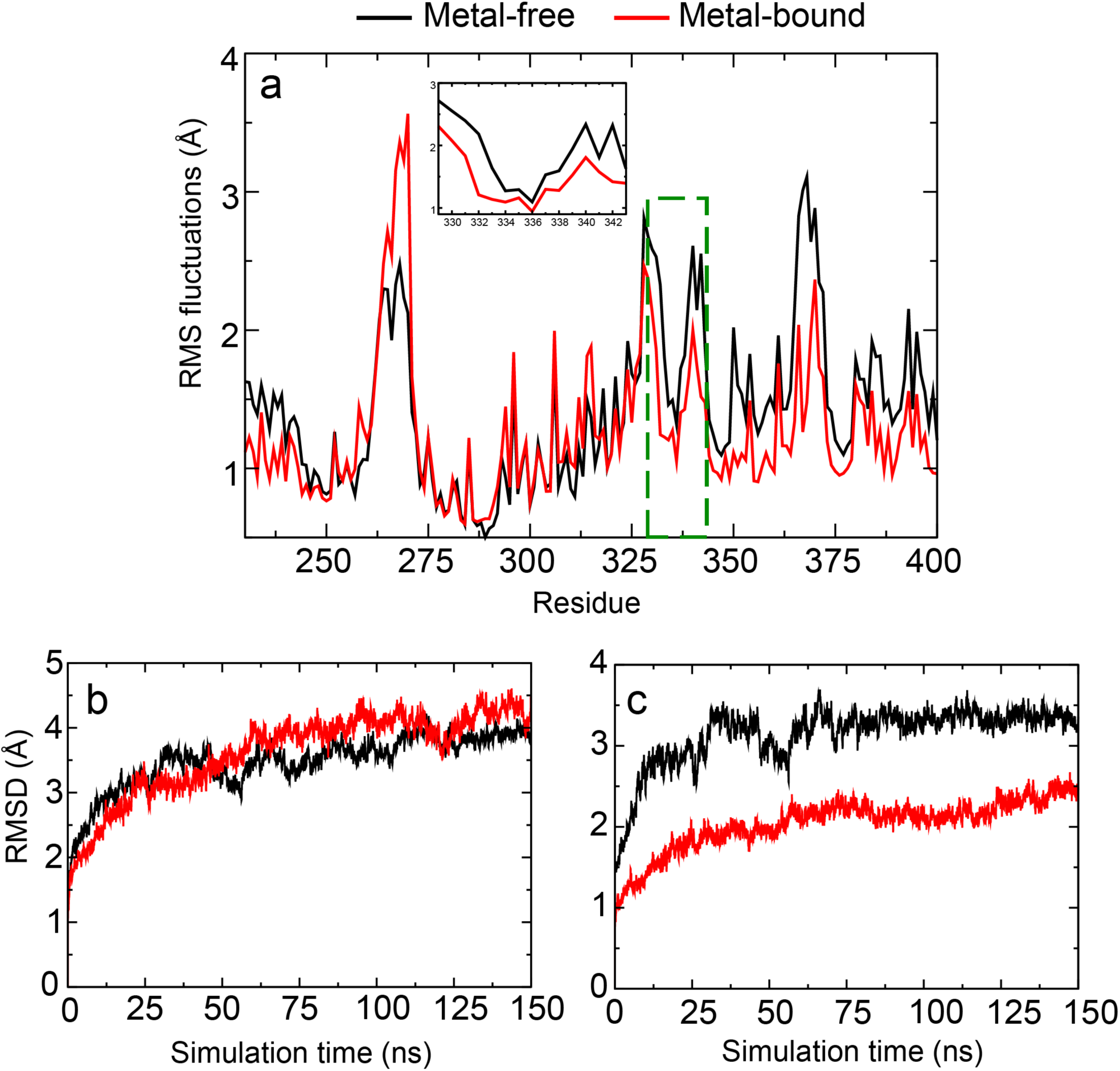
Structural and dynamic properties of hSlo1 obtained from simulations of the recently published Cryo-EM structures (PDB: 6V3G and 6V38). **a.** Root-Mean-Square Fluctuations (RMSF) profiles of residues 230 - 400 (PGD, C-linker and RCK1 N-lobe). C-linker RMSF is highlighted in the green dashed box and in insert. The Cα RMSD of **b.** the whole channel and **c.** residues 100 – 500 (the Core region) as a function of simulation time. RMSF and RMSD were obtained from the 150ns MD simulations with time step 0.05ns.

**Figure S3.**
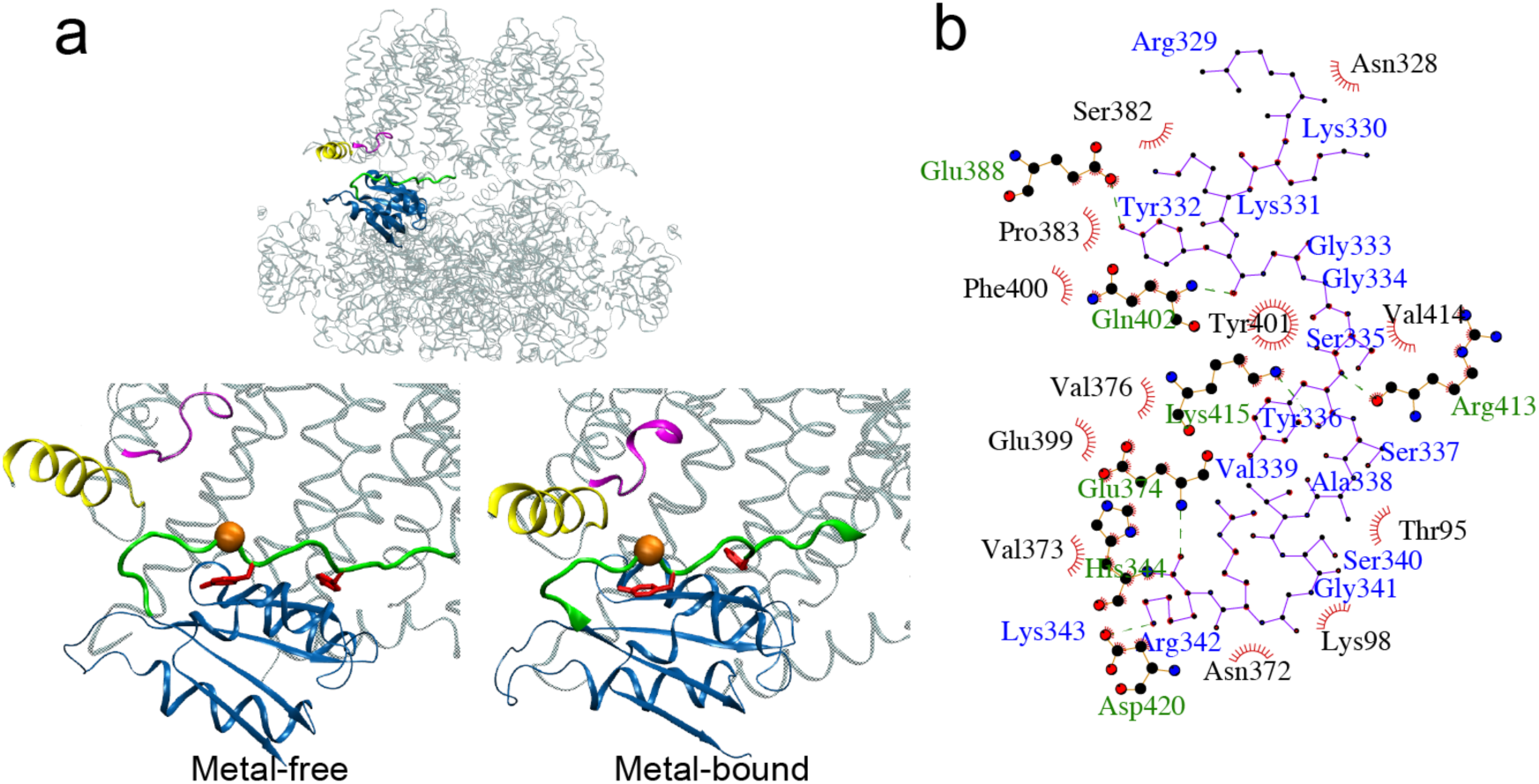
Orientation and packing between C-linker and VSD/RCK1 N-lobe of WT hSlo1. **a.** Segments highlighted are: C-linker (residue 329-343, green); RCK1 N-lobe (dark blue, residue 344-427); VSD S0’ (yellow, residue 92-107) and S2-S3 loop (magenta, residue 171-177). The orange sphere marks the position of S337 Cα; Y332 and Y336 are shown as red Licorice. **b.** Residue contact map of C-linker (blue labels) with VSD and RCK1 N-lobe (green and black labels) derived from the Cryo-EM structure of the metal-bound state of hSlo1.

**Figure S5.**
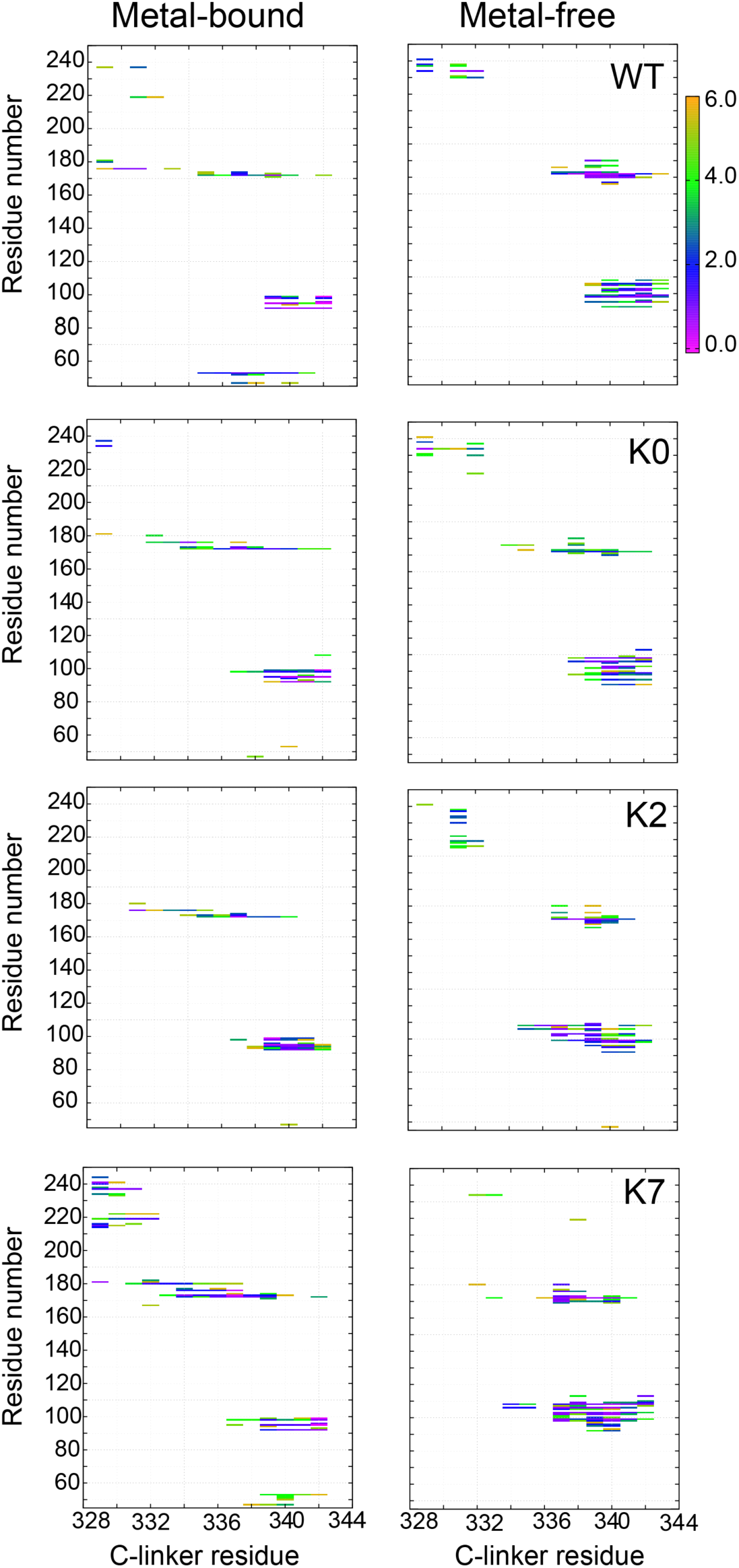
Probabilities of residue-residue contacts between the C-linker and VSD of the neighboring chains. VSD spans residues 45 – 250, where residues 45 to 53 belong to S0 helix, 92-107 to S0’, 171 - 177 to the S2-S3 loop, and 225 - 230 to the S4-S5 loop. The contact probabilities were calculated from atomistic simulations. See Methods for additional details.

**Figure S6.**
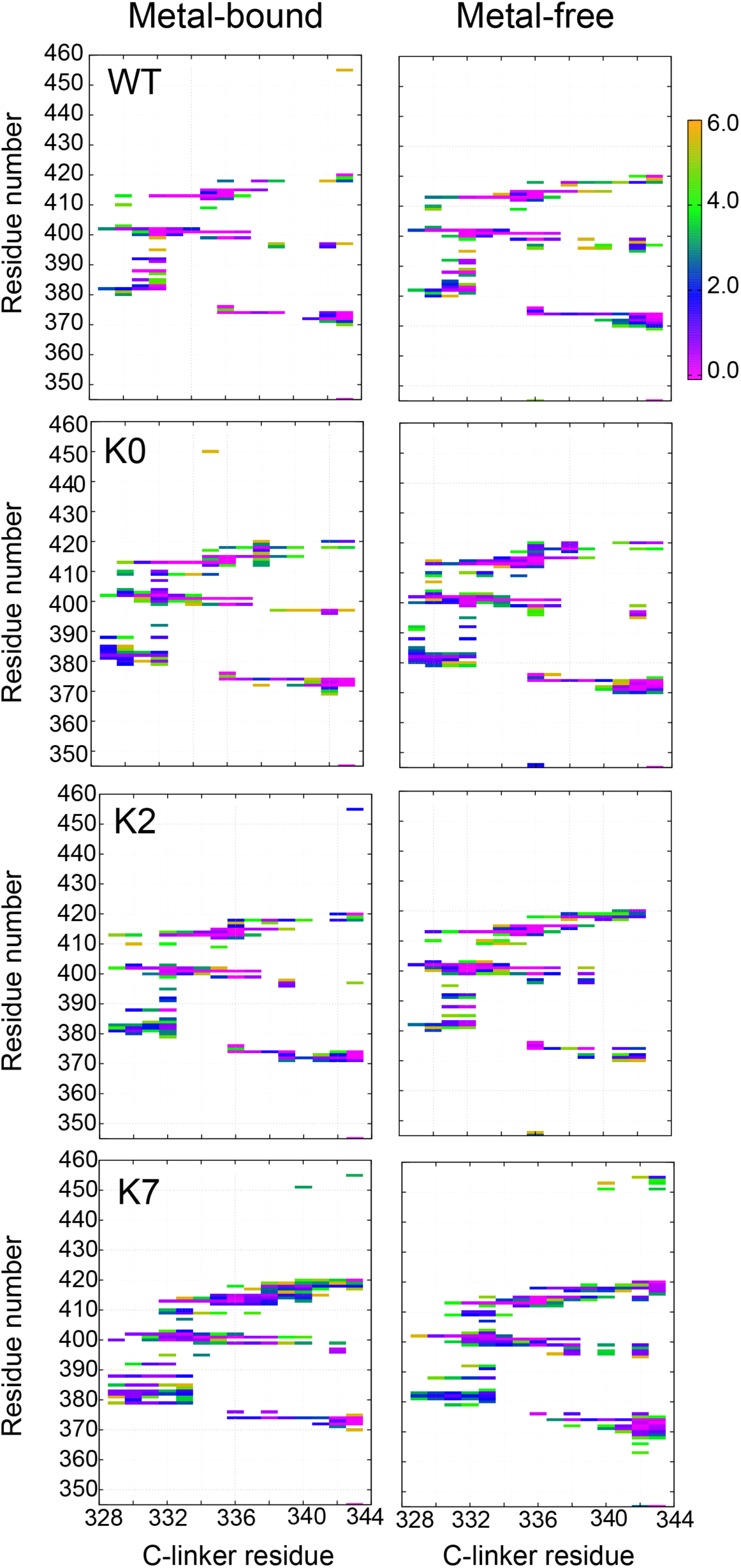
Probabilities of residue-residue contacts between the C-linker and RCK1 N-lobe of the same chain. See Methods for additional details.

**Figure S7.**
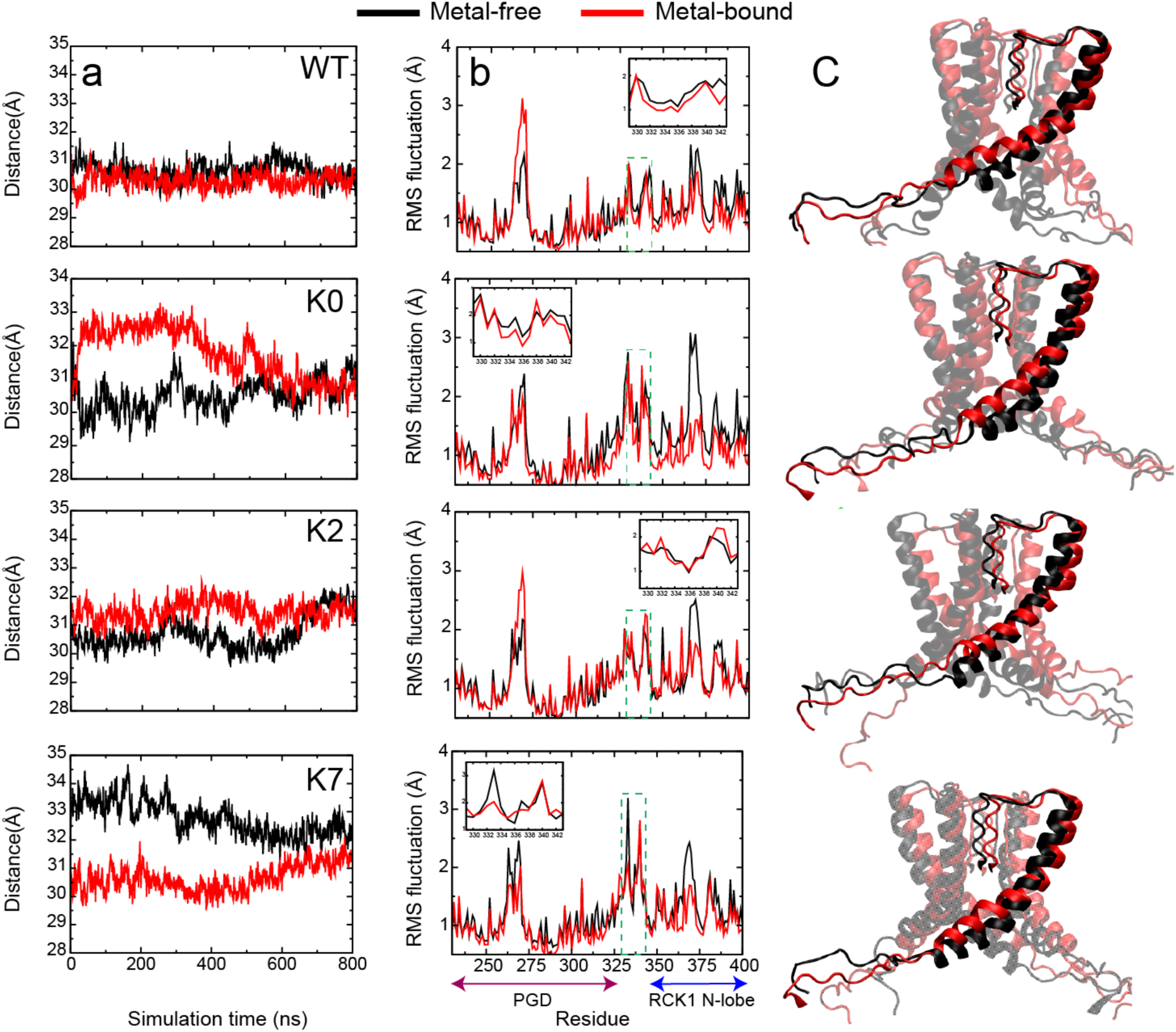
Structural and dynamic properties of the C-linker in WT hSlo1 and mutants. **a.** The C-linker N-C distance (329 – 343 Cα) as a function of time during 800-ns MD simulations. **b.** RMSF profiles of residues 230 - 400 (PGD, C-linker and RCK1 N-lobe) derived from the same MD trajectories, showing limited flexibility of the C-linker (green dashed box and in inserts). **c.** Representative snapshots showing the equilibrated conformations of C-linkers in both metal-bound and free states. See Methods for the details of the calculations.

**Figure S8.**
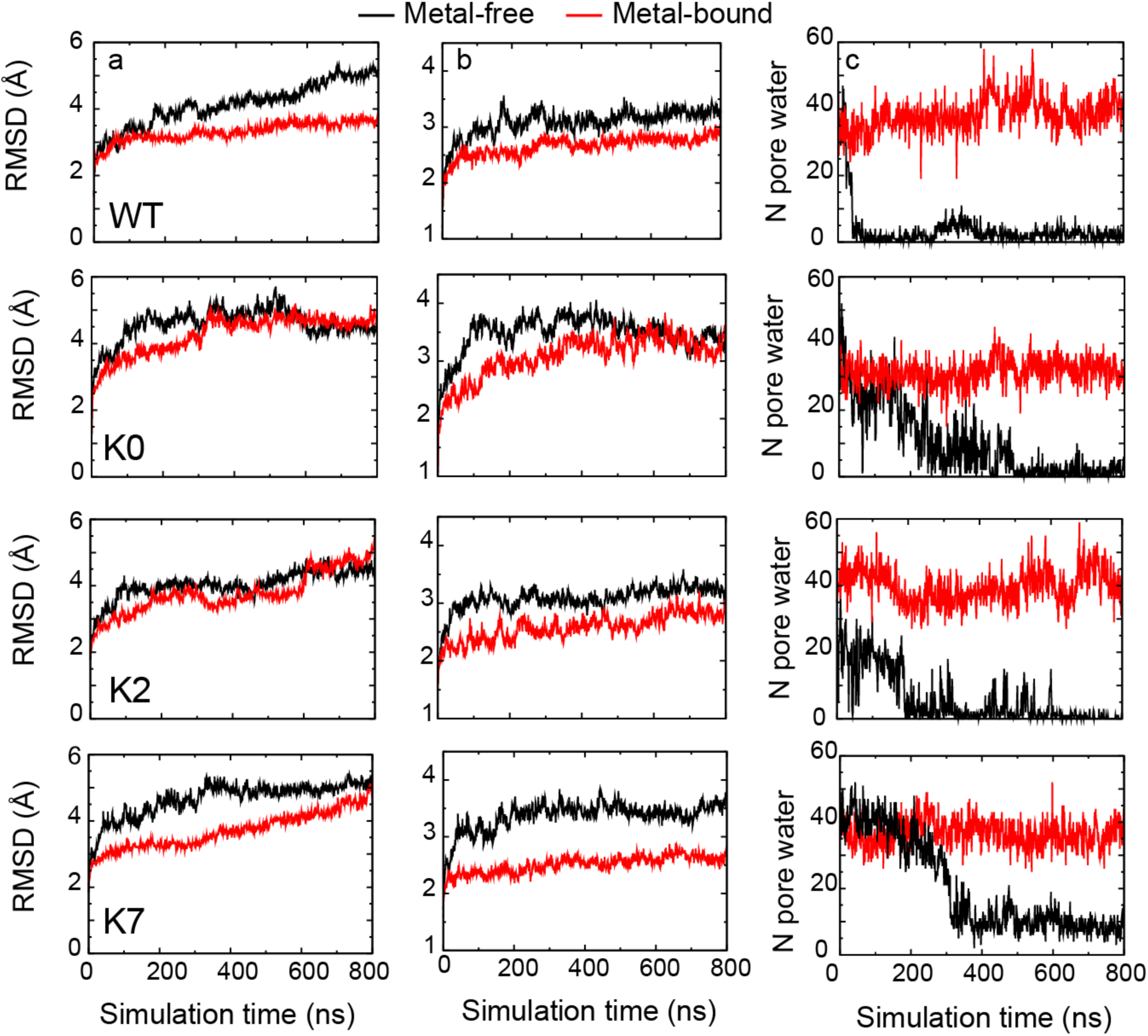
Structural and dynamic properties of WT hSlo1 and C-linker mutants. The Cα RMSD of **a.** the whole channel and **b.** residues 100 – 500 (the Core region) as a function of time during 800-ns MD simulations. **c.** Evolution of the number of pore water, showing that all hSlo1 constructs readily undergo dewetting transitions in the metal-free state. See Methods for additional details.

**Figure S9.**
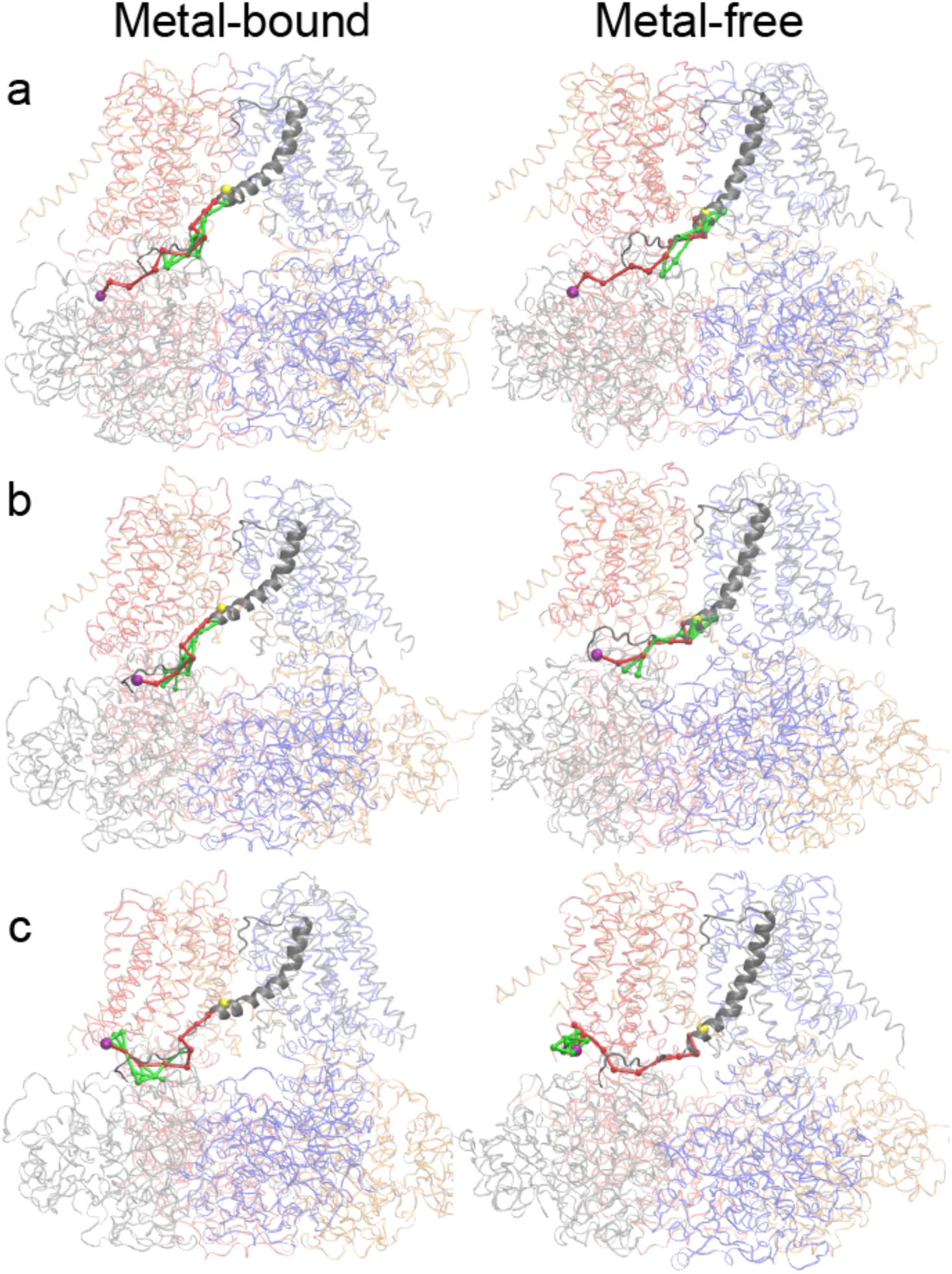
Optimal (red) and suboptimal (green) dynamic coupling pathways between the critical residues in the Ca^2+^ and Mg^2+^ binding sites (Cα colored as purple sphere) and I323 (Cα colored as yellow sphere) in PGD. Whole channel is shown as transparent ribbon with each chain colored differently. S6 and C-linker of chain C are colored in gray and shown with a cartoon representation. **a.** Path between R514 (located in RCK1 Ca^2+^ binding site in CTD; purple sphere) in chain C to I323 (located in S6 helix of PGD; yellow sphere) in chain C. **b.** Path from E374 (located in Mg^2+^ binding site in CTD; purple sphere) in chain C to I323 (located in S6 helix of PGD; yellow sphere) in chain C. **c.** Path from D99 (located in Mg^2+^ binding site in VSD; purple sphere) in chain B to I323 (located in S6 helix of PGD; yellow sphere) in chain C. See Methods for additional details.

**Figure S10.**
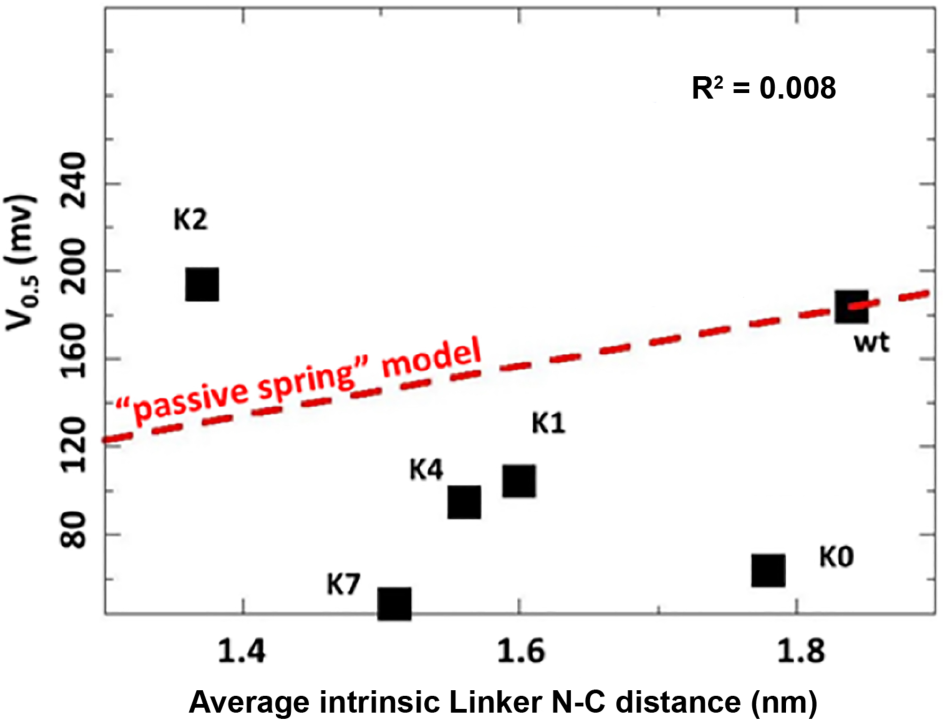
Correlation between the average intrinsic end-to-end distance of free C-linkers and measured V_0.5_ of the WT hSlo1 and mutants. The average distances were calculated from ABSINTH simulations of isolated C-linker peptides. The red dashed line shows the linear relationship derived from the C-linker insertion/deletion study (Niu et al 2004; also see Table S1).

**Figure S11.**
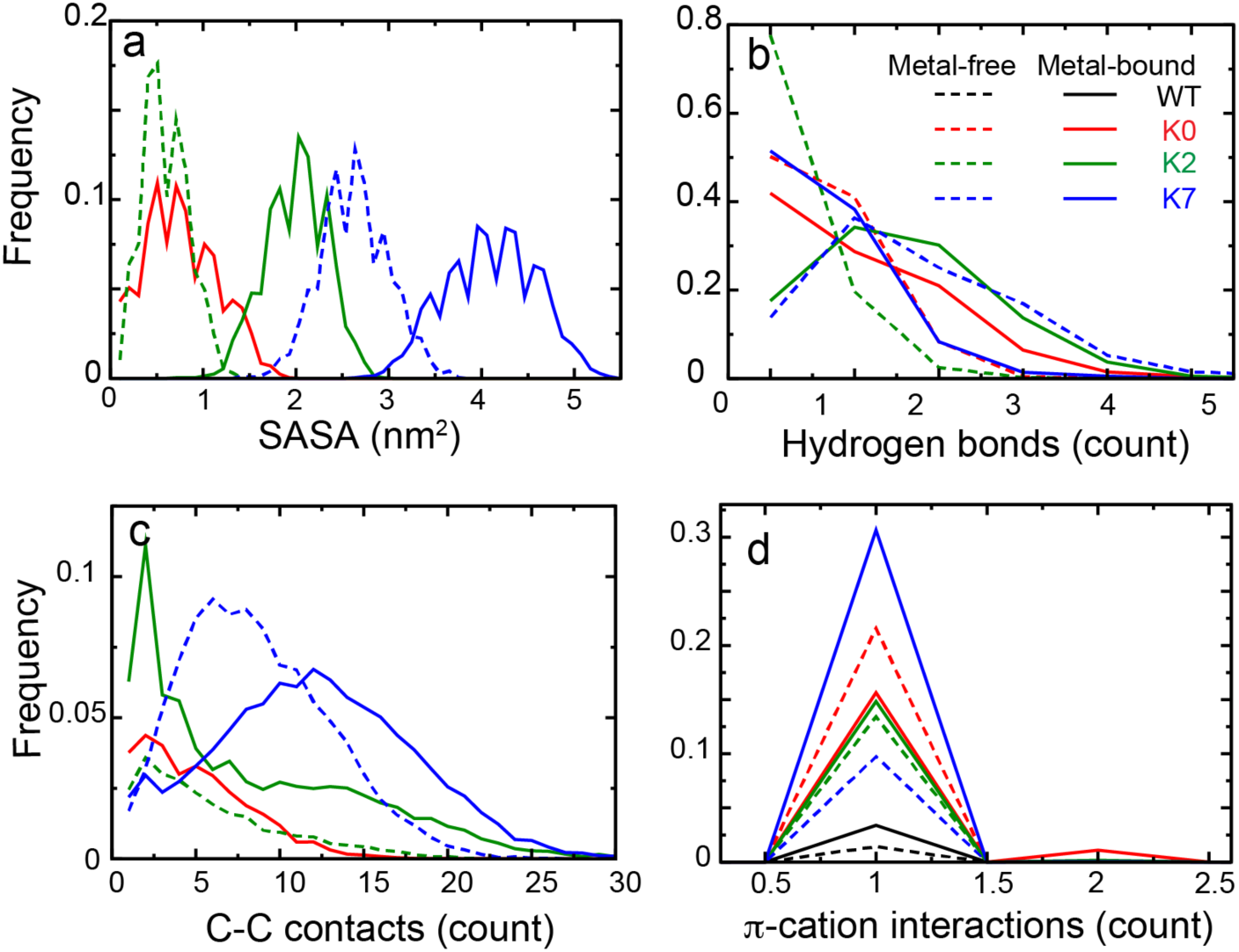
Distributions of various interactions of the C-linker Tyr residue nearest to the S6 helix, with the membrane interface for WT hSlo1 and mutants derived from simulations **a.** Tyr side chain SASA (Solvent Accessible Surface Area) of burial by lipid tails. **b.** Hydrogen bonding of Tyr OH with the lipid polar head groups. **c.** Carbon-carbon contacts between the Tyr aromatic ring and lipid tails. **d.** π-cation interactions between Tyr aromatic ring and POPC choline groups. Distributions derived from simulations of metal-free and bound states are shown in dashed and solid lines, respectively. No hydrophobic, hydrogen bonding, or C-C contacts as defined above were observed in the WT channel. See Methods for the details of the calculations.

**Figure S12.**
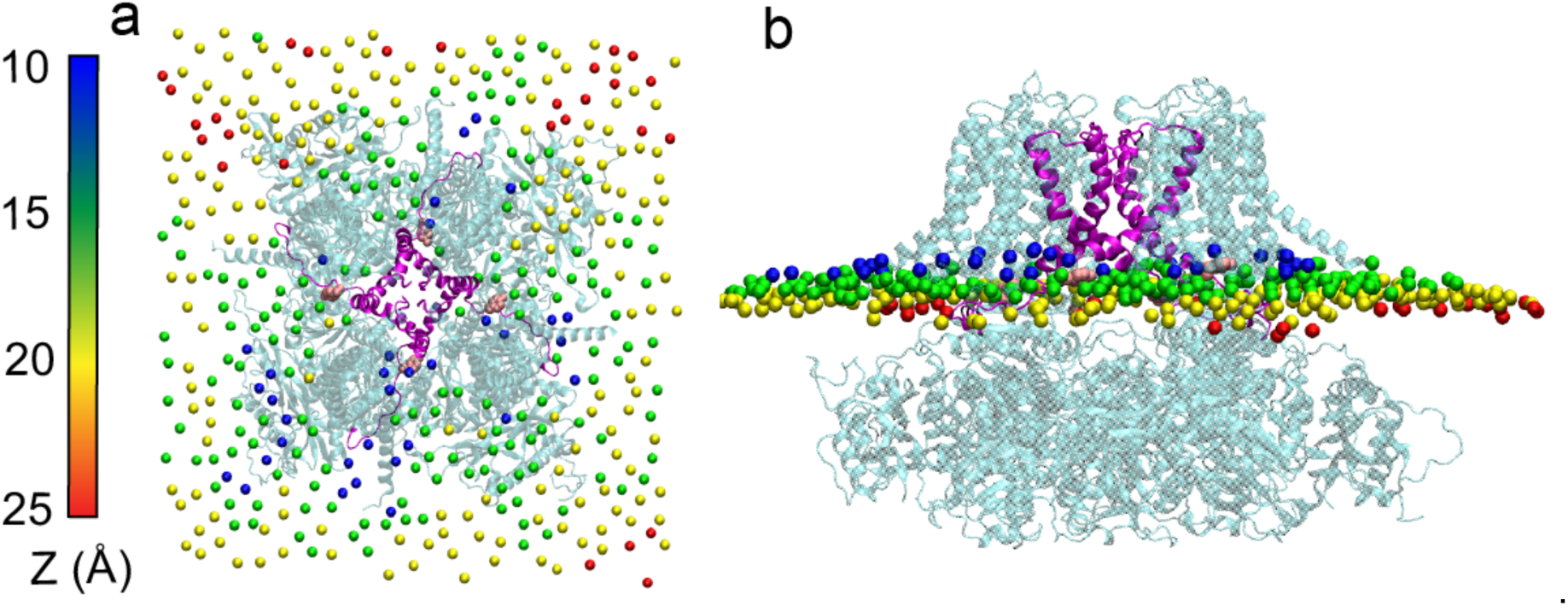
A representative snapshot of metal-bound K0 channel showing membrane distortion around the protein. **a.** Top view, **b.** side view. The POPC phosphorous atoms have been colored according to their distances to the membrane center. S6 helices are colored in magenta and Tyr 330 side chains are shown as pink spheres.

**Figure S13.**
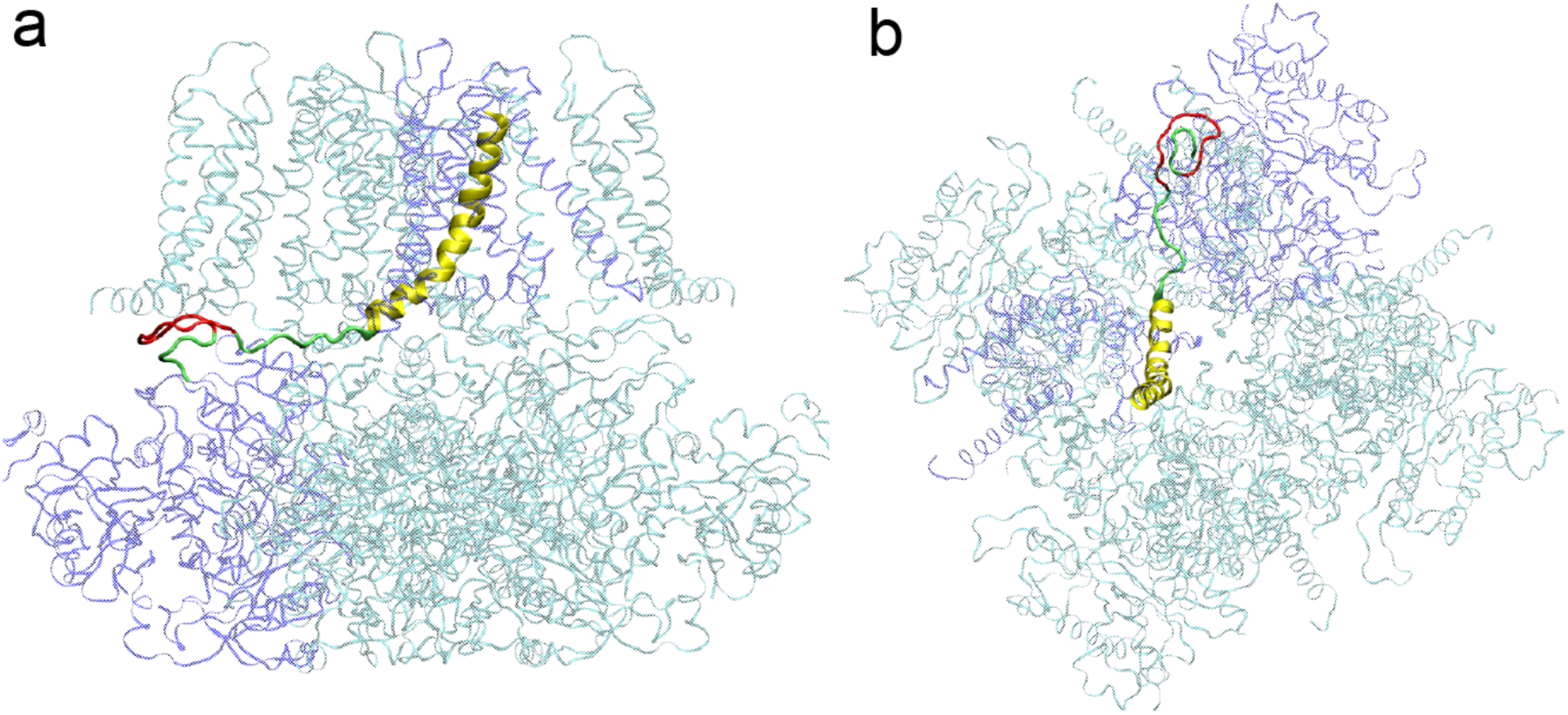
A representative structural model of hSlo1 with 12 residues (3x AAG units; colored red) inserted after S337 in the C-linker region (colored green). **a.** side view; **b.** top view. For clarity, only the C-linker and S6 helix (yellow) of chain A (colored as blue ribbon) are highlighted.

## References

1. Yang H, Zhang G, & Cui J (2015) BK channels: multiple sensors, one activation gate. Front Physiol 6:29.

2. Salkoff L, Butler A, Ferreira G, Santi C, & Wei A (2006) High-conductance potassium channels of the SLO family. Nat Rev Neurosci 7(12):921–931.

3. Knaus H, et al. (1996) Distribution of high-conductance Ca(^2+^)-activated K^+^ channels in rat brain: targeting to axons and nerve terminals. The Journal of Neuroscience 16(3):955–963.

4. Lee US & Cui J (2010) BK channel activation: structural and functional insights. Trends Neurosci. 33(9):415–423.

5. Uebele VN, et al. (2000) Cloning and functional expression of two families of β-subunits of the large conductance calcium-activated K^+^ channel. Journal of Biological Chemistry 275(30):23211–23218.

6. Horrigan FT & Aldrich RW (2002) Coupling between Voltage Sensor Activation, Ca^2+^ Binding and Channel Opening in Large Conductance (BK) Potassium Channels. The Journal of General Physiology 120(3):267–305.

7. Shi J & Cui J (2001) Intracellular Mg^2+^ Enhances the Function of Bk-Type Ca^2+^-Activated K^+^ Channels. The Journal of General Physiology 118(5):589–606.

8. Gessner G, et al. (2012) Molecular mechanism of pharmacological activation of BK channels. Proceedings of the National Academy of Sciences 109(9):3552–3557.

9. Zhou Y, Yang H, Cui J, & Lingle CJ (2017) Threading the biophysics of mammalian Slo1 channels onto structures of an invertebrate Slo1 channel. J. Gen. Physiol. 149(11):985–1007.

10. Yang HH, et al. (2008) Activation of Slo1 BK channels by Mg2+ coordinated between the voltage sensor and RCK1 domains. Nat. Struct. Mol. Biol. 15(11):1152–1159.

11. Tao X, Hite RK, & MacKinnon R (2017) Cryo-EM structure of the open high-conductance Ca2+-activated K+ channel. Nature 541(7635):46–51.

12. Hite RK, Tao X, & MacKinnon R (2017) Structural basis for gating the high-conductance Ca2+-activated K+ channel. Nature 541(7635):52–57.

13. Niu X, Qian X, & Magleby KL (2004) Linker-gating ring complex as passive spring and Ca(2+)-dependent machine for a voltage- and Ca(2+)-activated potassium channel. Neuron 42(5):745–756.

14. Jia Z, Yazdani M, Zhang G, Cui J, & Chen J (2018) Hydrophobic gating in BK channels. Nat. Commun. 9(1):3408.

15. Wilkens CM & Aldrich RW (2006) State-independent block of BK channels by an intracellular quaternary ammonium. J. Gen. Physiol. 128(3):347–364.

16. Schewe M, et al. (2019) A pharmacological master key mechanism that unlocks the selectivity filter gate in K+ channels. Science 363(6429):875-+.

17. Kopec W, Rothberg BS, & de Groot BL (2019) Molecular mechanism of a potassium channel gating through activation gate-selectivity filter coupling. Nat. Commun. 10(1):5366.

18. Tian Y, Heinemann SH, & Hoshi T (2019) Large-conductance Ca2+ - and voltage-gated K+ channels form and break interactions with membrane lipids during each gating cycle. Proc. Natl. Acad. Sci. U. S. A. 116(17):8591–8596.

19. Budelli G, Geng Y, Butler A, Magleby KL, & Salkoff L (2013) Properties of Slo1 K^+^ channels with and without the gating ring. Proceedings of the National Academy of Sciences 110(41):16657–16662.

20. Zhang G, et al. (2017) Deletion of cytosolic gating ring decreases gate and voltage sensor coupling in BK channels. The Journal of General Physiology 149(3):373–387.

21. Tao X & MacKinnon R (2019) Molecular structures of the human Slo1 K+ channel in complex with β4. eLife 8:e51409.

22. Eargle J & Luthey-Schulten Z (2012) NetworkView: 3D display and analysis of protein.RNA interaction networks. Bioinformatics 28(22):3000–3001.

23. LeVine MV & Weinstein H (2014) NbIT--a new information theory-based analysis of allosteric mechanisms reveals residues that underlie function in the leucine transporter LeuT. PLoS Comput. Biol. 10(5):e1003603.

24. McClendon CL, Kornev AP, Gilson MK, & Taylor SS (2014) Dynamic architecture of a protein kinase. Proc. Natl. Acad. Sci. U. S. A. 111(43):E4623–4631.

25. Das RK & Pappu RV (2013) Conformations of intrinsically disordered proteins are influenced by linear sequence distributions of oppositely charged residues. Proc. Natl. Acad. Sci. U. S. A. 110(33):13392–13397.

26. Vitalis A & Pappu RV (2009) ABSINTH: A New Continuum Solvation Model for Simulations of Polypeptides in Aqueous Solutions. J. Comput. Chem. 30(5):673–699.

27. Mao AH, Lyle N, & Pappu RV (2013) Describing sequence-ensemble relationships for intrinsically disordered proteins. Biochem. J. 449(2):307–318.

28. Monticelli L, et al. (2008) The MARTINI coarse-grained force field: Extension to proteins. J. Chem. Theory Comput. 4(5):819–834.

29. Johansson ACV & Lindahl E (2006) Amino-Acid Solvation Structure in Transmembrane Helices from Molecular Dynamics Simulations. Biophysical Journal 91(12):4450–4463.

30. Roberts MF, Khan HM, Goldstein R, Reuter N, & Gershenson A (2018) Search and Subvert: Minimalist Bacterial Phosphatidylinositol-Specific Phospholipase C Enzymes. Chem. Rev. 118(18):8435–8473.

31. Neale C, Hsu JC, Yip CM, & Pomes R (2014) Indolicidin binding induces thinning of a lipid bilayer. Biophys. J. 106(8):L29–31.

32. Budelli G, Geng YY, Butler A, Magleby KL, & Salkoff L (2013) Properties of Slo1 K+ channels with and without the gating ring. Proc. Natl. Acad. Sci. U. S. A. 110(41):16657–16662.

33. Hille B (2001) Ion Channels of Excitable Membranes (Oxford University Press, Sinauer, Sunderland, MA) 3rd Ed.

34. Magleby KL (2003) Gating mechanism of BK (Slo1) channels: So near, yet so far. The Journal of General Physiology 121(2):81–96.

35. Bavro VN, et al. (2012) Structure of a KirBac potassium channel with an open bundle crossing indicates a mechanism of channel gating. Nat. Struct. Mol. Biol. 19(2):158–163.

36. Šali A & Blundell TL (1993) Comparative protein modelling by satisfaction of spatial restraints. Journal of Molecular Biology 234(3):779–815.

37. Brooks BR, et al. (2009) CHARMM: The Biomolecular Simulation Program. J. Comput. Chem. 30(10):1545–1614.

38. Lee J, et al. (2016) CHARMM-GUI input generator for NAMD, GROMACS, AMBER, OpenMM, and CHARMM/OpenMM simulations using the CHARMM36 additive force field. Journal of Chemical Theory and Computation 12(1):405–413.

39. Jorgensen WL, Chandrasekhar J, Madura JD, Impey RW, & Klein ML (1983) Comparison of simple potential functions for simulating liquid water. The Journal of Chemical Physics 79(2):926–935.

40. Huang J, et al. (2017) CHARMM36m: an improved force field for folded and intrinsically disordered proteins. Nat. Methods 14(1):71–73.

41. D.A. Case, et al. (2014) AMBER 14 (University of California, San Francisco).

42. Darden T, York D, & Pedersen L (1993) Particle mesh Ewald: An *N*-log (*N*) method for Ewald sums in large systems. The Journal of Chemical Physics 98:10089.

43. Ryckaert J-P, Ciccotti G, & Berendsen HJC (1977) Numerical integration of the cartesian equations of motion of a system with constraints: molecular dynamics of n-alkanes. J. Comput. Phys. 23(3):327–341.

44. Chow K-H & Ferguson DM (1995) Isothermal-isobaric molecular dynamics simulations with Monte Carlo volume sampling. Computer Physics Communications 91(1):283–289.

45. Åqvist J, Wennerström P, Nervall M, Bjelic S, & Brandsdal BO (2004) Molecular dynamics simulations of water and biomolecules with a Monte Carlo constant pressure algorithm. Chemical Physics Letters 384(4):288–294.

46. Michaud-Agrawal N, Denning EJ, Woolf TB, & Beckstein O (2011) MDAnalysis: A toolkit for the analysis of molecular dynamics simulations. Journal of Computational Chemistry 32(10):2319–2327.

47. Hess B, Kutzner C, van der Spoel D, & Lindahl E (2008) GROMACS 4: algorithms for highly efficient, load-balanced, and scalable molecular simulation. J. Chem. Theory Comput. 4(3):435–447.

48. Abraham MJ, et al. (2015) GROMACS: High performance molecular simulations through multi-level parallelism from laptops to supercomputers. SoftwareX 1–2:19–25.

49. Humphrey W, Dalke A, & Schulten K (1996) VMD: Visual molecular dynamics. J. Mol. Graph. 14(1):33-&.

50. Glykos NM (2006) Software news and updates carma: A molecular dynamics analysis program. Journal of Computational Chemistry 27(14):1765–1768.

51. Floyd RW (1962) Algorithm 97: Shortest path. Commun. ACM 5(6):345.

52. Butler A, Tsunoda S, McCobb D, Wei A, & Salkoff L (1993) mSlo, a complex mouse gene encoding “maxi” calcium-activated potassium channels. Science 261(5118):221–224.

53. Shi J, et al. (2002) Mechanism of magnesium activation of calcium-activated potassium channels. Nature 418(6900):876–880.

